# The Emergence of the Metabolic Signaling of the Nucleoredoxin-like Genes during Evolution

**DOI:** 10.1101/2022.01.06.475223

**Authors:** Najate Aït-Ali, Frédéric Blond, Emmanuelle Clérin, Ala Morshedian, Quénol Cesar, François Delalande, Mitsumasa Koyanagi, Catherine Birck, John Han, Xiaoyuan Ren, Alain van Dorsselaer, Akihisa Terakita, Gordon L. Fain, Thierry Léveillard

## Abstract

The nucleoredoxin-like genes *NXNL1* and *NXNL2* were identified through the biological activity of rod-derived cone viability factors (RdCVF and RdCVF2), the alternatively spliced variants produced by intron retention, that mediate signaling between rod and cone photoreceptors by stimulating glucose uptake. These therapeutic genes for inherited retinal degenerations also produce by splicing thioredoxin-like proteins that reduce oxidized cysteines in photoreceptor proteins. The first *NXNL* genes date from the first animal phyla. Intron retention produces an active RdCVF protein in the tentacles of *Hydra vulgaris*, a species without eyes. A Scallop RdCVF protein is produced by ciliated photoreceptors of the retina and binds its receptor, BSG1. In the lamprey, a descendent of early vertebrates, RdCVF metabolic signaling between rod and cones is fully established. In the mouse, the production of BSG1 by photoreceptors is regulated by cell-specific splicing inhibition. RdCVF signaling predates photoreceptors and evolved through two alternative splicing events.

## INTRODUCTION

The *Nxnl1* gene was identified by high-content screening of a retinal expression library on a cone-enriched culture system through the activity of its product with intron retention, the rod-derived cone viability factor (RdCVF) (Leveillard et al., 2004; Millet-Puel et al., 2021). Splicing of this unique intron produces the thioredoxin-related protein RdCVFL, which protects photoreceptors against damage caused by oxidation of cysteines in proteins, such as the microtubule-associated protein τ (TAU) (Cronin et al., 2010; Elachouri et al., 2015; Fridlich et al., 2009). Intron retention is rod-specific as rods express both RdCVF and RdCVFL, while cones express only RdCVFL (Mei et al., 2016). Extracellular RdCVF binds the basigin-1 (BSG1)/GLUT1 (SLC2A1) complex at the surface of cones, which increases glucose uptake and aerobic glycolysis flux required for the daily renewal of photoreceptor outer segments (Ait-Ali et al., 2015; Chinchore et al., 2017). The enzymatic activity of the thioredoxin RdCVFL is supported by the metabolism of glucose through the pentose phosphate pathway, which is regulated in cones by RdCVF produced by rods (Leveillard and Ait-Ali, 2017). RdCVF and RdCVFL prevent the secondary cone degeneration of models of retinitis pigmentosa, the most frequent form of inherited retinal degeneration (Byrne et al., 2015; Mei *et al*., 2016). The metabolic and redox signaling of the *NXNL1* gene products is a mutation-independent treatment currently under development for that untreatable disease (Camacho et al., 2019; Clerin et al., 2020; Leveillard and Sahel, 2017).

The second member of the *NXNL* family, *NXNL2*, shares the same genic organization and produces a trophic factor, RdCVF2, by intron retention. RdCVF2 is a thioredoxin-like protein that inhibits TAU phosphorylation induced by light-damage (Chalmel et al., 2007; Elachouri et al., 2015). The *Nxnl2*^−/−^ mouse displays a progressive dysfunction of vision and olfaction (Jaillard et al., 2012). In the mouse, the main expression site of *Nxnl2* outside the eye is the *area postrema*, a circumventricular organ composed of fenestrated capillaries at the interface of the cerebrospinal fluid and blood circulation (Léveillard et al., 2021). The *Nxnl2*^−/−^ mouse has a memory deficit which progresses to a tauopathy during aging, reminiscent of the two phases of Alzheimer’s disease progression (Jaillard et al., 2021).

Intron retention of RdCVF mRNA leads to a disruption of the thioredoxin fold of RdCVFL, which implies that RdCVFL precedes RdCVF in the most simplistic evolutionary scenario. To trace the origin of RdCVF, we studied *NXNL* gene function in an early metazoan animal of the Cnidarian phylum, *Hydra vulgaris*; in the bivalve Mollusca, *Patinopecten yessoensis* (scallop); in an early cordate, *Branchiostoma lanceolatum* (amphioxus); in the jawless vertebrate, *Petromyzon marinus* (lamprey); and in a mammal, *Mus musculus* (mouse). Our study reveals that RdCVF metabolic signaling results from two successive alternative splicing events during evolution, first in the gene encoding the ligand, and then in that of the receptor, which together are at the origin of the rod-to-cone interaction.

## RESULTS

### Hydra RdCVFLa and RdCVFLb are enzymatically active thioredoxins

The *Hydra vulgaris* body is a hollow tube divided into two parts, the foot and the head (**Figure 1A**). The head is composed of the hypostome (mouth) at the apex and a zone where tentacles emerge from a ring. The *Hydra vulgaris* genome carries two tandemly arranged *NXNL* genes, *NXNLa* and *NXNLb* (**Figure 2A**), encoding for putative RdCVFLa and RdCVFLb proteins (**Figure S1**), whose mRNAs are expressed by the hydra polyp (**Figure 1B**). Their expression is ubiquitous, like the mitochondrial thioredoxin 2 *TXN2*, as shown by *in situ* hybridization with an antisense probe combining RdCVFLa/b (**Figure 1C**). RT-PCR confirms that both RdCVFLa and RdCVFLb mRNAs are expressed evenly by the hydra’s head and foot (**Figure 1D**).

**Figure 1.**
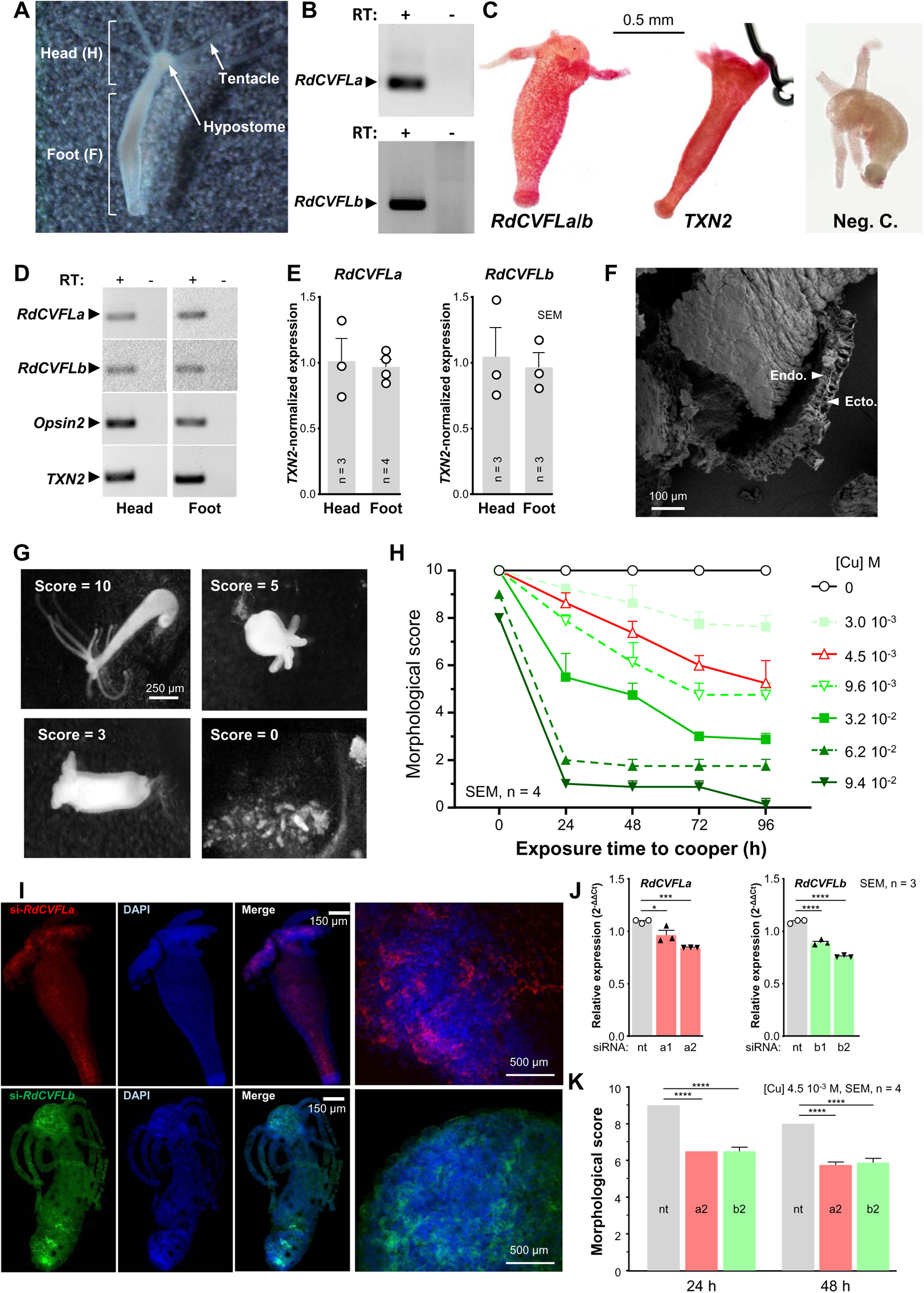
Hydra RdCVFLa and RdCVFLb are enzymatically active thioredoxins. (**A**) Hydra vulgaris morphology. (**B**) Expression of RdCVFLa and RdCVFLb mRNAs by hydra polyp. RT: reverse transcriptase. (**C**) *In situ* hybridization of RdCVFLa/b (the riboprobe is not discriminative), and TXN2. Neg. C.: negative control, the RdCVFLa/b sense riboprobe. (**D**) Expression of RdCVFLa, RdCVFLb, Opsin2 and TXN2 mRNAs in the head and foot of *Hydra vulgaris*. RT: reverse transcriptase. (**E**) Quantification of expression of RdCVFLa, RdCVFLb mRNAs in the head and the foot of hydra after normalization to TXN2 mRNA. (**F**) Scanning electron microscopy of hydra body showing the proximity of ectoderm (Ecto.) and endoderm (Endo.). (**G**) Representative images of hydra morphology scores. (**H**) Morphology scores of hydra after exposure to increasing concentration of copper. (**I**) Representative images of the delivery of si-RdCVFLa (red) and si-RdCVFLb (green) to the hydra body. Nuclei are labeled with DAPI. (**J**) Quantitative analysis of RdCVFLa and RdCVFLb mRNA expression after delivery of si-RdCVFLa a1 or a2 or si-RdCVFLb b1 or b2 as compared to non-targeting (nt) control, normalized to TXN2 mRNA. The data were analyzed with one-way ANOVA. (**K**) morphology scores of hydra polyps in the presence of 4.5 µM copper. The data were analyzed with one-way ANOVA.

**Figure 2.**
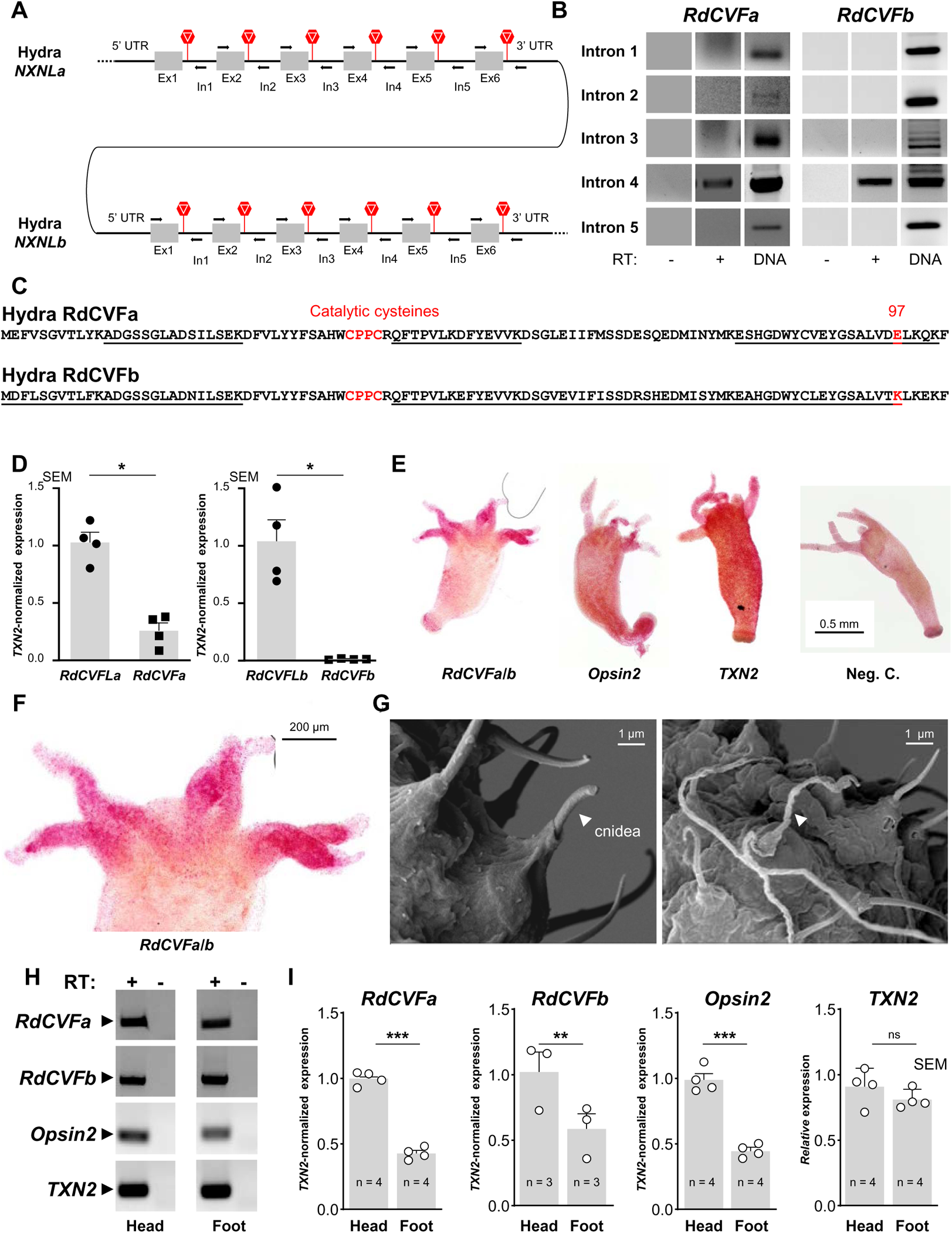
Hydra RdCVFa is expressed by cells of the tentacles. **(A**) The two tandemly repeated hydra *NXNLa* and *NXNLb* genes. Stop codons are represented by stop signs. Ex: exon, In: intron. (**B**) Analysis of the retention of the five introns of *NXNLa* and *NXNLb*. RT: reverse transcriptase. (**C**) Sequence of the hydra RdCVFa and RdCVFb proteins. The underlined sequences have been identified as tryptic peptides by mass spectrometry. The catalytic sites and the non-conserved E/K position 97 are indicated in red. (**D**) Quantification of expression of RdCVFa and RdCVFb mRNAs in the hydra body after normalization to TXN2 mRNA. (**E**) *In situ* hybridization of RdCVFa/b (the riboprobe is not discriminative) and TXN2. Neg. C.: negative control, the RdCVFa/b sense riboprobe. (**F**) Higher magnification of the signal of the RdCVFa/b antisense riboprobe in the head of hydra. (**G**) Scanning electron microscopy of hydra tentacle showing the cnidea (white arrowhead), before and after cnidocyte discharge. (**H**) Expression of RdCVFa, RdCVFb, Opsin2 and TXN2 mRNAs in the head and the foot of *Hydra vulgaris*. RT: reverse transcriptase. (**I**) Quantification of expression of RdCVFLa, RdCVFLb, and Opsin2 mRNAs in the head and the foot of hydra after normalization to TXN2 mRNA. Relative expression of TXN2 mRNA.

*Cnidarians* such as *Hydra vulgaris* have no mesoderm; the ectoderm and the endoderm are in direct contact (**Figure 1F**). External molecules reach all body surfaces directly in this simple diploblastic organism. Copper added in the medium containing a hydra produces reactive oxygen species (ROS) by the Fenton-like reaction (Schumann et al., 2002). ROS preferentially oxidizes thiols in proteins, triggering tissue damage leading to morphological changes of hydra that are classified by means of decimal scores (**Figure 1G**). Score 10 corresponds to a hydra with extended tentacles, score 5 to a contracted hydra with tentacles, score 3 to an expanded hydra with shorter tentacles, and 0 to a disintegrated hydra (Wilby et al., 1990). Using this scoring system, we evaluated the effect of increasing the concentration of copper on hydra morphology and show the dependence of toxicity on copper concentration and time (**Figure 2H**), in accordance with previous observations (Karntanut and Pascoe, 2000). Oxidized cysteines are reduced by thioredoxins such as RdCVFL (Ren et al., 2017). Fluorescent siRNAs (Tortiglione et al., 2009) targeting hydra RdCVFLa (a1 and a2) or RdCVFLb (b1 and b2) transduce cells of the entire hydra body (**Figure 1I**), which results in a significant downregulation of their mRNAs (**Figure 1J**). The toxicity of 4.5 µM Cu is exacerbated by reducing the expression of both RdCVFLa and RdCVFLb (**Figure 1K**). These observations demonstrate that RdCVFLa and RdCVFLb are enzymatically active thioredoxins involved in the defense of the whole organism against oxidative damage.

### Hydra RdCVFa is expressed by cells of the tentacles

We designed primers within the 5 introns of the *NXNLa* and *NXNLb* genes to test for intron retention (**Figure 2A**). Each of these introns has a stop codon in-frame with the coding sequence of their respective 5’ exons. RT-PCR shows that intron retention occurs only for intron 4, for both genes (**Figure 2B**). These alternatively spliced mRNAs encode for two putative proteins of 104 residues, RdCVFa and RdCVFb (**Figure 2C**). We identified tryptic peptides of these two proteins by proteomic analysis of the hydra body, confirming their existence. Still, the peptides do not discriminate between thioredoxins (RdCVFLa and RdCVFLb) and truncated thioredoxins (RdCVFa and RdCVFb). Interestingly, these two hydra proteins correspond closely to biologically active RdCVF (Leveillard *et al*., 2004) (**Figure S2A**). The two retained introns 4 have no sequence conservation and do not significantly differ from the others in terms of secondary structure, 3’ to the donor sites (**Figure S2B**), and to the content of pyrimidine or thymidine (%TC: 62.4% ±6) in the pyrimidine tract, 5’ to the acceptor sites (**Figure S2C**).

In the whole hydra body, RdCVFa and RdCVFb mRNAs represent ~30% and 0.005% of RdCVFLa and RdCVFLb mRNAs, respectively (**Figure 2D**). We compared the expression of the mRNAs resulting from intron retention to that of TXN2, a ubiquitous mitochondrial thioredoxin; and OPSIN2, an opsin expressed by photosensitive cells of *Hydra vulgaris* (**Figure 2E**). We observed a dense, punctuated labeling of the RdCVFa/b probe in the tentacles (**Figure 2F**). Despite a relatively simple anatomical organization and the absence of an eye, *Hydra vulgaris* responds to light (Guertin and Kass-Simon, 2015; Passano and McCullough, 1962). It has photosensitive neurons which use ciliary photoreceptor transduction, localized on the surface of the hydra body and expressing *OPSIN2* (Plachetzki et al., 2012).

The phylum *Cnidaria* is defined by the presence of cnidocytes. Those cells contain one giant secretory organelle called a cnidocyst. Upon stimulation of sensory neurons, *Hydra vulgaris* cnidocytes eject cnidae (cnidocysts) containing poison to capture their prey (**Figure 2G**). We studied the expression of RdCVFa and RdCVFb mRNAs in the head relative to the foot by RT-PCR (**Figure 2H**). RdCVFa, RdCVFb and OPSIN2 mRNAs are expressed at a higher level in the head, while TXN2 mRNA is evenly expressed (**Figure 2I**). By similarity to OPSIN2, the regionalized expression of RdCVFa and RdCVFb in the head of *Hydra vulgaris* argues for a biological function.

### RdCVFa is the ligand of a hydra cell surface receptor

The truncated thioredoxin proteins RdCVFa and RdCVFb are most likely enzymatically inactive, like RdCVF in the mouse (Leveillard *et al*., 2004), but they are candidate ligands for a cell surface receptor homologous to BSG1 (Ait-Ali *et al*., 2015). Basigin is homologous to neuroplastin (NPTN) (Muramatsu and Miyauchi, 2003). We identified neural-cell-adhesion-molecule-like-1 (NCAM1) as the protein in the hydra proteome most closely related to the basigin / neuroplastin superfamily, even though NCAM1 does not contain the immunoglobulin (Ig) domain Ig0 which is required for its interaction with RdCVF (**Figure S3A**) (Ait-Ali *et al*., 2015).

We designed primers for this non-Ig domain to study its expression (**Figure 3A**). We observed NCAM1 expression by hydra polyps but did not detect an alternatively spliced mRNA that would correspond to BSG2, the chaperone of lactate transporters (**Figure 3B**) (Tokar et al., 2017). The predicted NCAM1 protein was validated by proteomic analysis of the hydra body (**Figure 3C**). We designed alkaline phosphatase (AP) fusion proteins to test for the interaction of RdCVFa and RdCVFb with NCAM1 at the surface of HEK293-transfected cells (**Figure 3D**). The putative ligands were checked by western blotting using both anti-AP and anti-GFP, the latter a linker in the construction (**Figure 3E**). The expression of NCAM1 and mouse BSG1 were validated using hemagglutinin (HA) epitope (**Figure 3F**). We did not detect any increase in the purple staining of HEK293 cells in the presence of NCAM1, indicating that RdCVFa and RdCVFb do not interact with NCAM1 under such conditions (**Figure 3G**).

**Figure 3.**
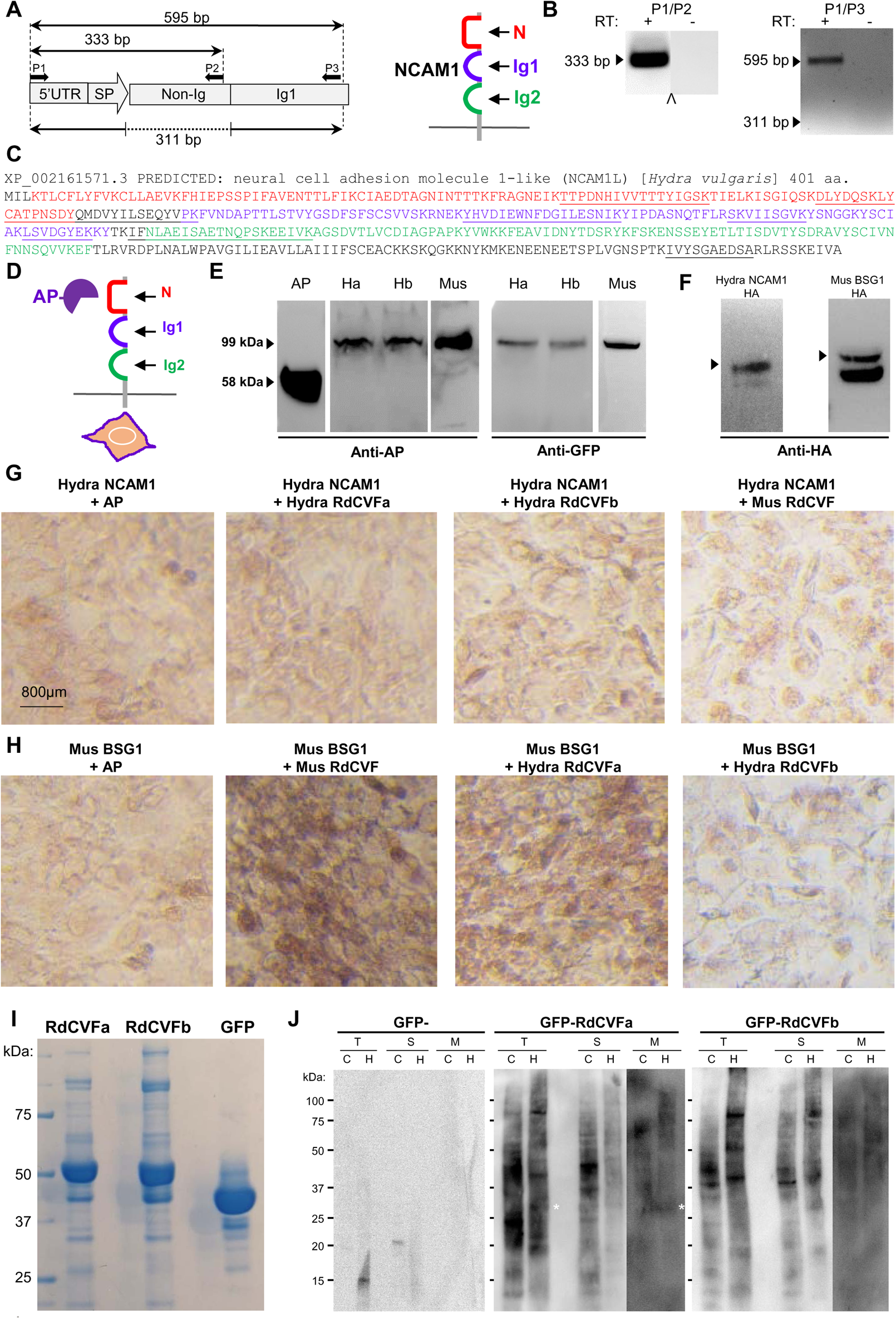
RdCVFa is the ligand of a hydra cell-surface receptor. (**A**) Schematic representation of the 5’ part of the NCAM1 mRNA and drawing of the NCAM1 protein in the membrane. The N domain is not homologous to an Ig domain. (**B**) Expression of NCAM1 mRNA (P1/P2 and P1/P3) and the absence of the shorter alternatively spliced form (P1/P3). (**C**) Amino-acid sequence of NCMA1 with the N-terminal domain (N), the Ig1, and the Ig2 domains colored in red, blue and green, respectively. The underlined sequences have been identified as tryptic peptides by mass spectrometry. (**D**) schematic representation of the alkaline-phosphatase binding assay. AP: alkaline phosphatase. (**E**) Analysis of the secretion of AP-fusion proteins after transfection of HEK293 cells. Ha and Hb: hydra AP-GFP-RdCVFa and AP-GFP-RdCVFb, Mus: mouse AP-GFP-RdCVF. (**F**) Analysis of the expression of HA-fusion proteins after transfection of HEK293 cells. (**G**) AP-binding assays. AP: unfused alkaline phosphatase. (**H**) AP-binding assays. (**I**) Coomassie staining gel of purified GFP-RdCVFa, GFP-RdCVFa, and GFP. (**J**) Far-western blotting analysis of total (T), cytoplasmic (S), and membranal (M) fractions of hydra body (H) and COS-1 cells (C). Specific signals detected by GFP-RdCVFa in the total and membranal fractions of *Hydra vulgaris* are indicated with white asterisks.

Mouse RdCVF interacts with mouse BSG1 but not with hydra NCAM1, in agreement with the participation of Ig0 in this interaction (Ait-Ali *et al*., 2015). Surprisingly, we found that hydra RdCVFa, but not RdCVFb, interacts with mouse BSG1 in this assay (**Figure 3H**). Such interaction validates hydra RdCVFa as a ligand for an unknown cell surface receptor, distinct from NCAM1. We used a far-western approach to look for RdCVFa and RdCVFb binding proteins. His-tagged RdCVFa and RdCVFb, fused to GFP, were produced in *E. coli* and purified by affinity chromatography (**Figure 3I**). Total (T), cytoplasmic (S), and membranal (M) fractions of the whole hydra body (H), and COS-1 cells (C) used as a negative control, were probed with the candidate ligands. We detected a specific signal at 25 kDa with GFP-RdCVFa but not with GFP-RdCVFb (**Figure 3J**). This signal is absent in the cytoplasmic fraction, and its intensity increases from the total to the membranal fraction, suggesting that the protein is expressed at the surface of hydra cells.

The analysis of the sequence of RdCVFa and RdCVFb points to an E to K transition at position 97 (**Figure 2C**), reminiscent of the E64K mutation identified in patients suffering from a form of Leber congenital amaurosis that disrupts the interaction with BSG1 (Ait-Ali *et al*., 2015; Hanein et al., 2006). In the structural model of RdCVFL proteins (Chalmel *et al*., 2007), this position is encompassed in an external α-helix (**Figure S3B**). We speculate that this transition K > E is the evolutionary origin of RdCVF signaling, due to the higher ratio of RdCVFa/RdCVFLa versus RdCVFb/RdCVFLb in *Hydra vulgaris* (**Figure 2D**). The absence of E at this conserved E/D position in the ctenophore proteome (**Figure S3C**) supports the view that only RdCVFa is biologically functional. The presence of an E at that position in *Spongilla Lacustris* RdCVFL is conflicting. Still, the absence of an intron in the *NXNL* gene of the sponge excludes intron retention and the production of an RdCVF protein by that species, as well for ctenophore *NXNLs*. *NXNL* is rather specifically expressed by the choanocyte, a sponge’s cell with a flagellum used to capture prey (Musser et al., 2021).

### RdCVF and RdCVFL are expressed by ciliated scallop photoreceptors

We next analyzed RdCVF signaling in the scallop *Patinopecten yessoensis*, a mollusc from the Pacific Northwest with up to 200 eyes lining the mantle tissue (**Figure 4A**) (Wang et al., 2017). The scallop *NXNL* gene has three coding exons (**Figure 4B**). In the scallop eye, we discovered that exons 1 and 2 are systematically spliced (P1/P4), and intron 1 is not retained (P1/P2). Intron 2 is alternatively spliced, giving an amplification product of 375 bp (*, P1/P5) or 192 bp (*, P3/P5), and a spliced mRNA (P1/P6 and P3/P6) (**Figure 4C**). This finding is like that in *Hydra vulgaris*. Intron 2 has a stop codon in-frame with the coding sequence of exon 2, which produces RdCVF and RdCVFL proteins of 107 and 141 residues (**Figures S1** and **S2**). The peculiarity of the scallop eye is the presence of both ciliary and rhabdomeric photoreceptors (Kojima et al., 1997). *Drosophila* eyes, like all insects, have only rhabdomeric photoreceptors and no *NXNL* gene (Chalmel *et al*., 2007; Funato and Miki, 2007). The ciliary and rhabdomeric photoreceptors of scallop are arranged in two separated retinas, a distal and a proximal retina (Palmer et al., 2017). We visualized the distal retina using anti-PScop2, a photopigment of ciliary photoreceptors (Arenas et al., 2018) (**Figure 4D** and **S4A**); for the proximal retina we used anti-Gqα (Kingston et al., 2017) (**Figure 4E** and **S4B**). In the scallop eye, both the RdCVFL and RdCVF mRNAs are specifically expressed by the distal retina containing ciliary photoreceptors (**Figure 4F**). *NXNL* expression is not restricted to the eye, since we also detected RdCVFL and RdCVF mRNAs in the gills, gonads, and muscle (**Figure 4G**).

**Figure 4.**
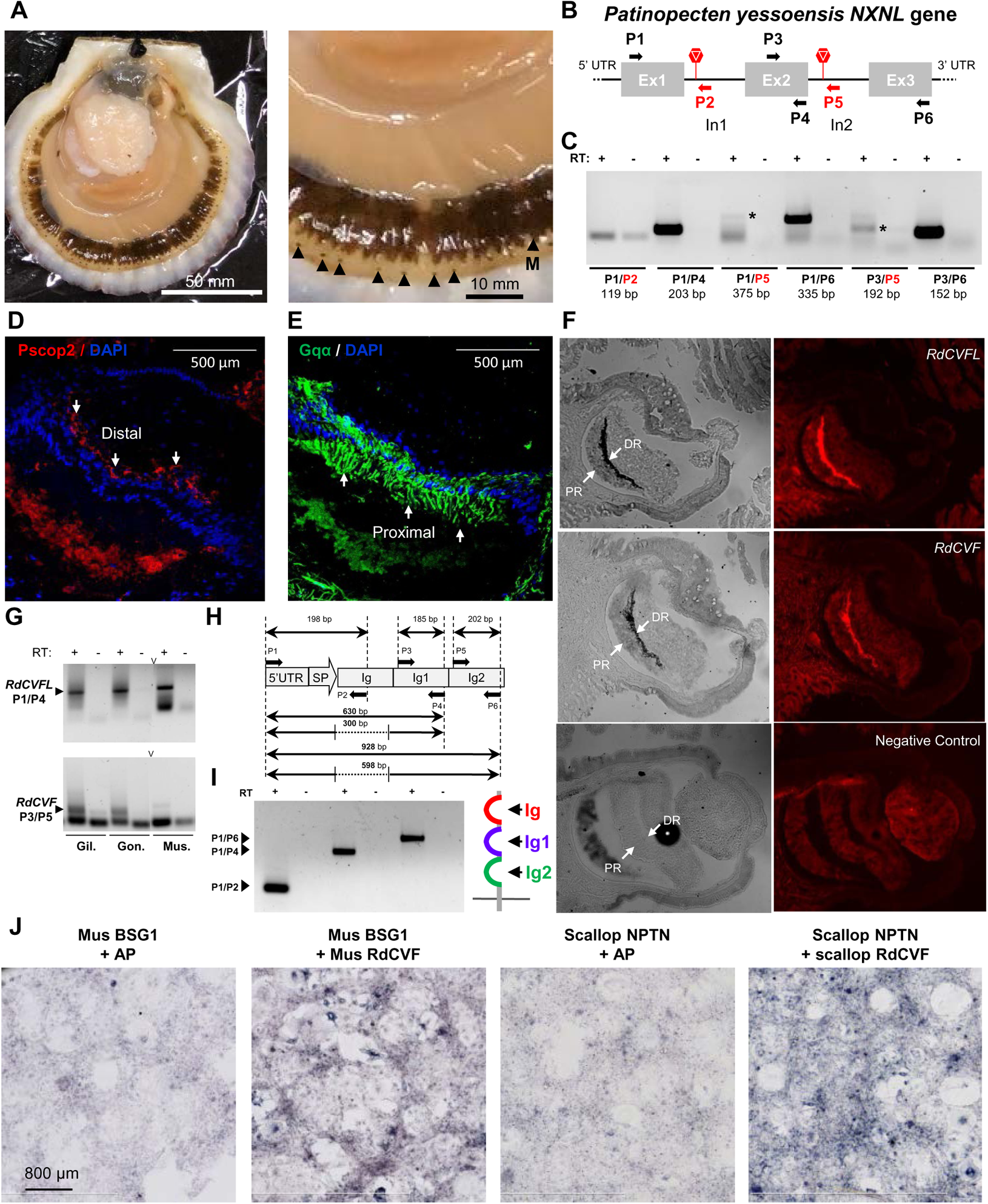
RdCVF and RdCVFL are expressed by ciliated scallop photoreceptors. (**A**) The Pacific scallop *Patinopecten yessoensis* and its multiple eyes are indicated by black arrowheads. M: mantle. (**B**) The *Patinopecten yessoensis NXNL* gene and its three coding exons. Stop codons are represented by stop signs. Ex: exon, In: intron. (**C**) Expression of RdCVFL and RdCVF in scallop eyes. RT: reverse transcriptase. (**D**) Expression of Pscop2 by the distal retina by immunohistochemistry. (**E**) Expression of Gqα protein by the proximal retina. (**F**) Expression of RdCVFL and RdCVF mRNAs in the distal retina (DR) by *in situ* hybridization. PR: proximal retina (**G**) Expression of RdCVF and RdCVFL mRNAs in scallop tissues; Gil.: gill, Gon.: gonad, Mus.: muscle. RT: reverse transcriptase. (**H**) Schematic representation of BSG1 mRNA. SP: signal peptide. (**I**) Expression of BSG1 by scallop eyes. RT: reverse transcriptase. (**J**) AP-binding assays.

The *Patinopecten yessoensis* proteome contains a neuroplastin-like protein of 396 residues (XP_021372255.1, NCBI) with three extracellular Ig domains (**Figure 4H**). In the scallop eye, we detected an mRNA encoding for a protein with three Ig domains, similar to BSG1, but we could detect no mRNA encoding for the splice variant proteins BSG2 without the N-terminal Ig domain. (**Figure 4I** and **Figure S4C**). To study the interaction of scallop RdCVF with BSG1, we produced AP-fusion proteins validated by western blotting (**Figure S4D**). Scallop RdCVF interacts with scallop BSG1, in the same way that mouse RdCVF interacts with mouse BSG1 (**Figure 4J**).

### Amphioxus expresses the putative *RdCVFa* receptor BSG1 and the chaperone BSG2

*Branchiostoma lanceolatum* (amphioxus) is an animal model of early vertebrate evolution (Escriva, 2018). Amphioxus has multiple photoreceptors: the frontal eye and lamellar body contain ciliary photoreceptors, whereas Joseph cells and dorsal ocelli have rhabdomeric photoreceptors (**Figure 5A**) (Lacalli, 2004; Suzuki et al., 2015). We identified four *NXNL* genes, arbitrarily named *NXNLa-NXNLd*, each with two exons in the *Branchiostoma lanceolatum* genome (Marletaz et al., 2018; Putnam et al., 2008).

**Figure 5.**
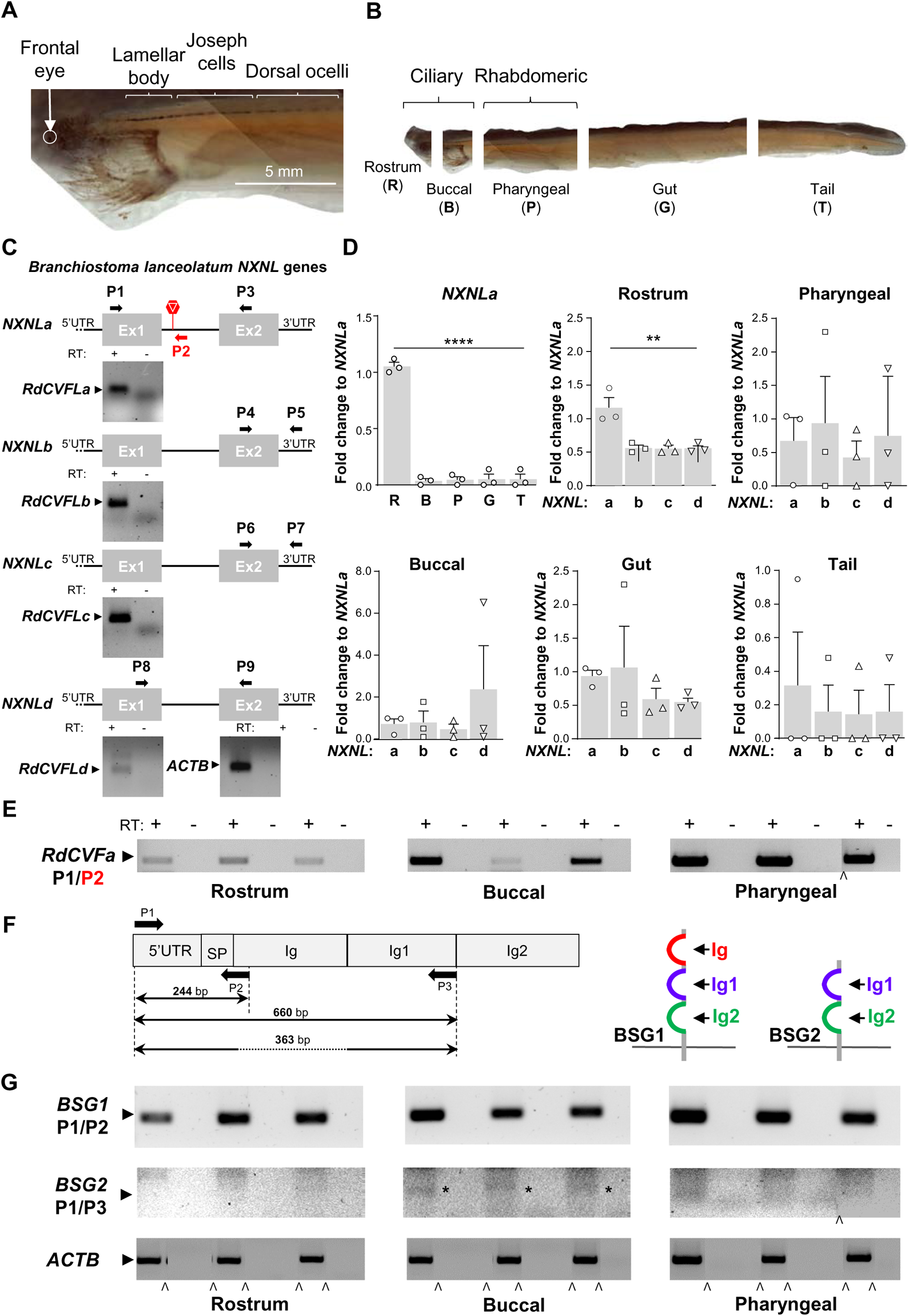
Amphioxus expresses the putative *RdCVFa* receptor BSG1 and the chaperone BSG2. (**A**) The head of amphioxus. (**B**) Sections of the amphioxus body used in expression studies. (**C**) The *Branchiostoma lanceolatum NXNL* genes *NXNLa* – *NXNLd* and their expression in the whole amphioxus body. The stop codon of *NXNLa* is represented by a stop sign. RT: reverse transcriptase. ACTB: cytoplasmic actin. (**D**) Relative expression of *NXNLa* mRNA by amphioxus body parts. Expression of *NXNLa*-*d* mRNA by amphioxus body parts normalized to that of *NXNLa* mRNA. (**E**) Expression of RdCVFa mRNA by three amphioxus body parts. (**F**) Schematic representation of BSG1 mRNA and drawing of the BSG1 and BSG2 proteins in the membrane. SP: signal peptide. (**G**) Expression of BSG1, BSG2, and ACTB mRNAs by three amphioxus body parts.

We sectioned the adult amphioxus body into five parts to study *NXNLs* expression (**Figure 5B**). The rostrum, where the frontal eye is located, was separated from the buccal part, that contains the lamellar body. RdCVFL mRNAs were detected in the whole amphioxus body (**Figure 5C** and **Figure S1**). The most striking observation is the restricted expression of *NXNLa* to the rostrum (**Figure 5D**). NXNLa is not expressed by Joseph cells and the dorsal ocelli, which are rhabdomeric, as NXNL for the scallop eye. None of the other *NXNL* genes shares a similar expression pattern. *NXNLb-d* genes are only marginally expressed in the amphioxus body apart from the rostrum, normalized by the *NXNLa* expression level.

The rostrum contains the frontal eye with ciliary photoreceptors, reminiscent of the regionalized expression of *NXNL* in the eye of the scallop (**Figure 4F**). The lamellar body reaches its greatest axial extent in late-stage larvae, when it spans most of the length of somite 1. It fragments after metamorphosis, leaving only scattered cells behind (Lacalli, 2004). The frontal eye has a neural connection to brain regions homologous with the caudal diencephalon and midbrain, which is also in the rostral part of the body. As a result, we could not firmly establish whether *NXNLa* is expressed by photoreceptors of the frontal eye (Lacalli, 2018). We observed the retention of the unique intron of the *NXNLa* gene leading to the expression of a putative amphioxus RdCVFa protein expressed by the rostral, buccal, and pharyngeal parts of the amphioxus using a non-quantitative assay (**Figure 5E** and **Figure S2A**).

The *Branchiostoma lanceolatum* genome also contains a gene homologous to *BSG*, encoding for a protein (BL11268_cuf1, Amphiencode) with three extracellular Ig domains similar to BSG1 (**Figure 5F** and **Figure S3A**). Its mRNA is distributed like that of RdCVFa (**Figure 5G**). We detected an mRNA encoding for BSG2 (Ochrietor et al., 2003). The amplified product was validated by sequencing the splicing junctions as for all RT-PCR products of our study (see STAR methods).

### The expression of RdCVF is restricted to rods of the lamprey retina

Hagfish and lamprey are cyclostomes and the only jawless vertebrates to have survived up to the present day (Fain, 2020). The genome of the lamprey (*Petromyzon marinus*) has a unique *NXNL* gene, annotated *NXNL1* in public databases, with two coding exons encoding for a RdCVFL protein after splicing (UniProt S4RDF3) (**Figure 6A** and **Figure S1**) (Smith et al., 2013). Intron retention in the lamprey retina produces an RdCVF mRNA in addition to that of RdCVFL (**Figure 6B** and **Figure S2A**). RdCVF expression represents one quarter of that of RdCVFL in the retina (**Figure 6C**). The expression of RdCVFL and RdCVF is restricted to the retina, as in the mouse (**Figure 6D**) (Reichman et al., 2010).

**Figure 6.**
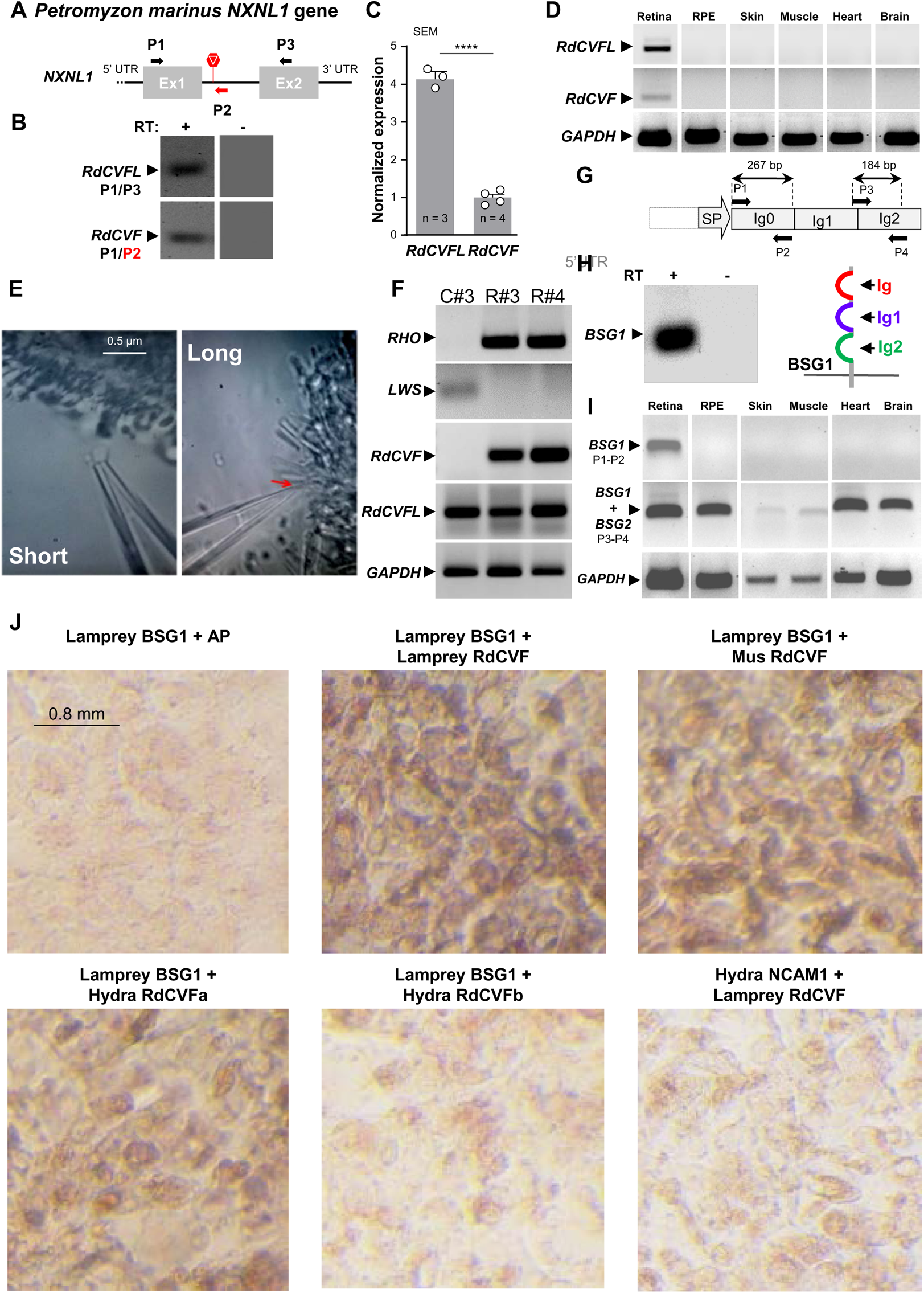
The expression of RdCVF is restricted to rods of the lamprey retina. (**A**) The *Petromyzon marinus NXNL1* gene. The stop codon is represented by a stop sign. (**B**) Expression of RdCVFL and RdCVF mRNAs in the lamprey retina. RT: reverse transcriptase. (**C**) Quantitative analysis of the expression of RdCVFL and RdCVF mRNA by lamprey retina. (**D**) Expression of RdCVFL, RdCVF, and glyceraldehyde-3-phospahte dehydrogenase (GAPDH) mRNAs by lamprey organs. RPE: retinal pigment epithelium. (**E**) Representative images of the collection by microaspiration with patch-clamp glass electrodes of a rod and a cone. The red arrow indicates the position of the electrodes and the long photoreceptor (cone). (**F**) Expression of RHO, long-wavelength cone opsin (LWS), RdCVF, RdCVFL, and GAPDH mRNAs by isolated cones and rods. (**H**) Expression of BSG1 by the lamprey retina and drawing of the BSG1 protein in the membrane. (**I**) Expression of BSG1, BSG1 + BSG2, and GAPDH mRNAs by lamprey organs. (**J**) AP-binding assays.

Rods evolved from cones, and progenitors of lamprey had rods and diverged from other vertebrates before the emergence of disks and the characteristic rod outer-segment ultrastructure (Morshedian and Fain, 2015). Lamprey rods and cones respond to light much like rods and cones in mammals (Morshedian and Fain, 2017). By microaspiration with a patch-clamp glass electrode, we collected rods (short, **Video S1**) and cones (long, **Video S2**) (**Figure 6E**). Photoreceptor identity, and the absence of contamination by material from surrounding cells, were validated by amplifying *RHO* and long-wavelength sensitive (*LWS*) cone opsin in rod and cone preparations (**Figure 6F**). Rods express both RdCVFL and RdCVF mRNAs, while cones express only RdCVFL mRNA as in the mouse retina (Mei *et al*., 2016).

The lamprey *BSG* gene encodes for a BSG1 protein (S4RAQ8, UniProt) with three extracellular Ig domains (**Figure 6G** and **Figure S3A**). This protein was detected with mass spectrometry (**Figure S6A**). Since the assembly of the lamprey genome is incomplete, we could not use 5’UTR *BSG* sequences to analyze possible alternative splicing and the production of a *BSG2* mRNA directly (Timoshevskiy et al., 2019). Nevertheless, RT-PCR primers specific for BSG1 mRNA show its restricted expression in the retina, while primers that can amplify BSG1 + BSG2 show that BSG2 mRNA is more broadly expressed (**Figure 6H, I**). Because rods express RdCVF and the retina expresses BSG1, we tested their interaction using AP-fusion proteins (**Figure S6B**). Lamprey BSG1 interacts with lamprey and mouse RdCVF, as well as hydra RdCVFa; but it does not interact with hydra RdCVFb (**Figure 6J**). Nor does lamprey RdCVF interact with hydra NCAM1. Lamprey RdCVF signaling resembles that in mammals.

### RdCVF receptor expression results from splicing inhibition in the mouse eye

In the mouse retina, RdCVF is expressed specifically by rods; and BSG1, the splicing product of the *Bsg* gene, is expressed by both rods and cones (Ait-Ali *et al*., 2015; Mei *et al*., 2016). We constructed successively two mouse models of the specific disruption of BSG1. The *Bsg1*^−/−v1^ mouse has two in-frame stop codons in exon 2 (**Figure 7SA**), and the *Bsg1*^−/−v2^ mouse has one in-frame stop codon in a different position of exon 2 (**Figure 7SC**). We studied the visual phenotype of homozygous mice at postnatal day (PN) 40 (**Figure 7SA** and **Figure 7SD**). *Bsg1*^−/−v1^ mice have a reduced scotopic *a*-wave amplitude at PN40 compared to both *Bsg1*^+/+^ and *Bsg1*^−/−v2^ mice (**Figures 7A** and **7B**). All mice are on a pure C57BL/6J background, so that the dysfunction of rods does not result from the confounding *rd8* allele (Mattapallil et al., 2012). At 12 months (M) of age, rod function does not distinguish *Bsg1*^+/+^ from *Bsg1*^−/−v2^ mice (**Figure 7C**).

**Figure 7.**
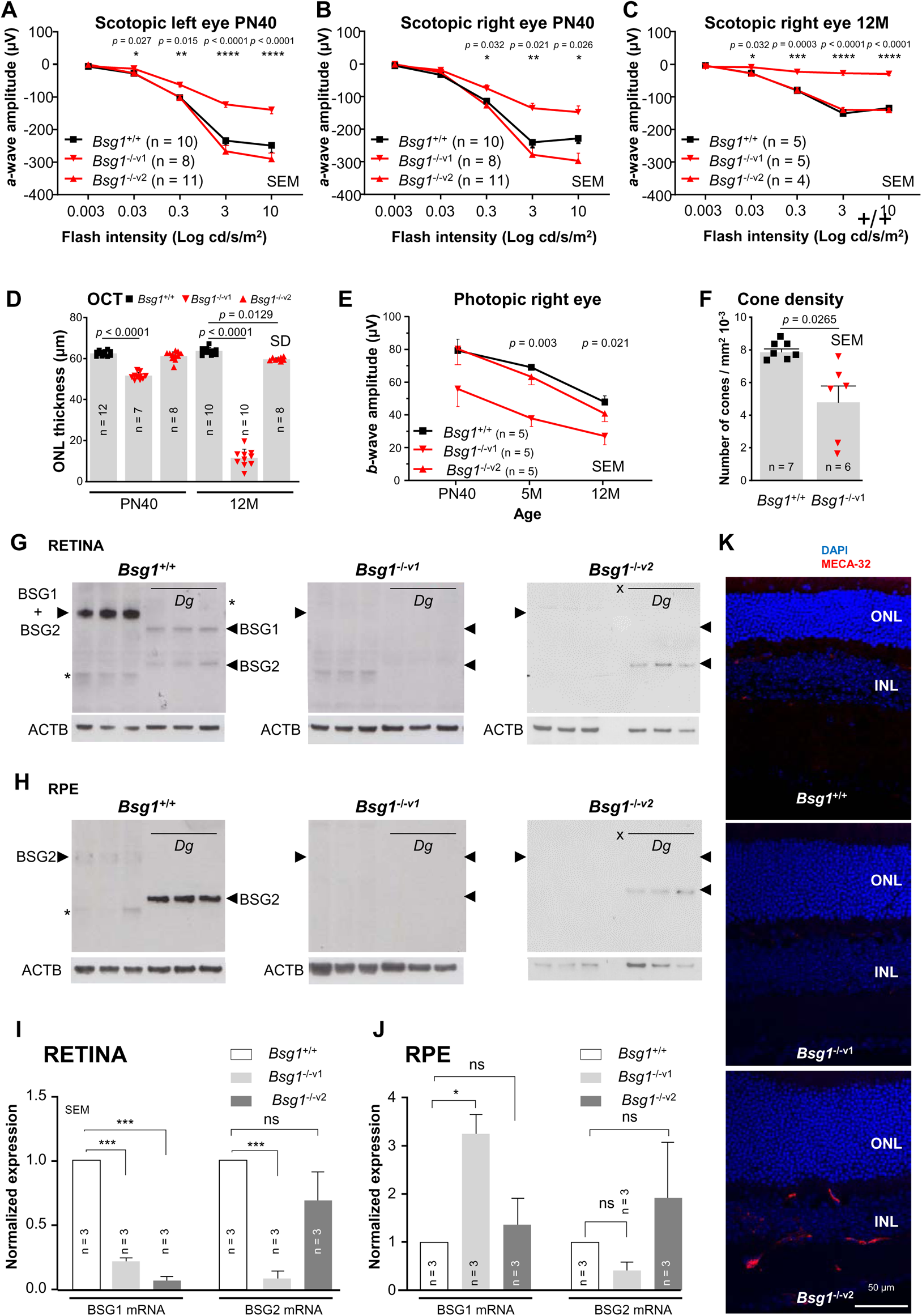
RdCVF receptor expression results from splicing inhibition in the mouse eye. (**A**) *a*-wave amplitude of the scotopic (rod-dependent) electroretinogram (ERG) of left eyes of the *Bsg* mutant mice at post-natal day (PN) days 40. (**B**) *a*-wave amplitude of the scotopic ERG of right eyes of *Bsg* mutant mice at PN40. (**C**) *a*-wave amplitude of the scotopic ERG of right eyes of the mutant mice at 12 months (M). (**D**) Thickness of the outer nuclear layer (ONL) of *Bsg* mutant mice at PN40 and 12M measured by optical coherence tomography (OCT). SD: standard deviation. (**E**) *b*-wave amplitude of the photopic (cone-dependent) ERG of right eyes of the *Bsg* mutant mice at P40, 5 and 12 months. (**F**) Cone density of *Bsg*^+/+^ and *Bsg*^−/−v1^ mutant retinas at 12M. (**G**) Expression of BSG1, BSG2, and ACTB by the retina of the *Bsg*^−/−v1^ mutant retina at PN40, before and after deglycosylation (*Dg*). The position of the BSG1 and BSG2 deglycosylated proteins is indicated by black arrowheads. ACTB: cytoplasmic actin, x: empty well. (**H**) Expression of BSG1, BSG2, and ACTB by the retinal pigment epithelium (RPE) of *Bsg*^−^ mutant mice at PN40, before and after deglycosylation (*Dg*). (**I**) Quantification of expression of BSG1 and BSG2 mRNAs by the retina *Bsg*^−^ mutant mice at PN40. (**J**) Quantification of expression of BSG1 and BSG2 mRNAs by the RPE of *Bsg*^−/−^ mutant mice at PN40. The data were analyzed with one-way ANOVA.

To look at rod survival, we measured the thickness of the outer nuclear layer (ONL) using spectral-domain optical coherence tomography (SD-OCT). At PN40, the retina of the *Bsg1*^−/−v1^ mice is thinner than both *Bsg1*^+/+^ and *Bsg1*^−/−v2^ retinas (**Figure 7D**). The thickness of this layer is further reduced in *Bsg1*^−/−v1^ mice at 12M, showing that rods which make up 97% of the ONL, are dying progressively. *Bsg1*^−/−v1^ mice have a reduced photopic *b*-wave amplitude from PN40 compared to *Bsg1*^+/+^ and *Bsg1*^−/−v2^ mice (**Figure 7E**). The deficit in cone vision of *Bsg1*^−/−v1^ mice at 12M is correlated with reduced cone survival compared to *Bsg1*^+/+^ mice (**Figure 7F**) (Clerin et al., 2011).

We analyzed the results of homologous recombination by western blotting. Deglycosylation (*Dg*) of protein extracts was used to visualize the migration of BSG1 and BSG2 in retinal extracts (Ait-Ali *et al*., 2015; Bai et al., 2014) (**Figure 7G**). In the *Bsg1*^−/−v2^ retina, the mutation in exon 2 results in the loss of BSG1, while in the *Bsg1*^−/−v1^ retina both BSG1 and BSG2 are lost. As expected, the RPE of the *Bsg1*^−/−v2^ mouse expresses specifically BSG2, but that is not the case for the *Bsg1*^−/−v1^ mouse (**Figure 7H**) (Han et al., 2020). We quantified the expression of the mRNAs encoding these two *Bsg* isoforms by qRT-PCR. Similar to the western blot data, BSG1 mRNA is reduced in both *Bsg1*^−/−v1^ and *Bsg1*^−/−v2^ retinas, but the expression of BSG2 mRNA not containing the mutations is also reduced in the *Bsg1*^−/−v1^ retina (**Figure 7I**). In the RPE of the *Bsg1*^−/−v1^ mouse, on the other hand, the reduction of expression of the BSG2 mRNA is correlated with an increase in expression of BSG1 mRNA (**Figure 7J**). Since the exon contains two stop codons, this mRNA is not translated into a BSG1 protein.

We analyzed the sequence of exon 2 to look for possible effects of the introduction of the stops codons (**Figure S6E**). We realized that the binding element of HNRNP (P2, H1, H2, H3) and F are erased by the stop codon introduced 5’ of exon 2 in the *Bsg1*^−/−v1^ mouse (**Figure S6F**). In the *rd1* retina, an autosomal recessive model of retinitis pigmentosa, HNRNPF and HNRNPH2 mRNAs have an expression profile resembling that of rods and cones (Blond and Leveillard, 2019). Using western blotting, we confirmed that HNRNPF is preferentially expressed by the outer retina containing rods and cones (**Figure S6G**). The RPE also expresses HNRNPF. HNRNPH2 is expressed by the inner retina and by the RPE but little or not at all by photoreceptors in the outer retina of the adult mouse. This is consistent with a possible role of HNRNPH2 inhibiting the production of the BSG1 mRNA through binding to an element that is absent in the *Bsg1*^−/−v1^ mouse genome. The visual phenotype of the *Bsg1*^−/−v1^ mouse may result from lactate accumulation in the outer retina (Ochrietor and Linser, 2004), as observed after deletion of the *Bsg* gene in the RPE at 4 weeks of age (Han *et al*., 2020). In the RPE, BSG2 is a chaperone for the lactate transporters MCT1 and MCT3 which mediate transepithelial transport from the subretinal space to the blood circulation of lactate, produced by the retina through aerobic glycolysis (Klipfel et al., 2021; Leveillard et al., 2019).

We observed a very small yet significant decrease in the ONL of the *Bsg1*^−/−v2^ mice at 12 months of age, which suggests that there are either changes to the rods or cones, which could explain the slight reduction in the *b*-wave of these animals at 12 months of age. Because the *Bsg1*^−/−v2^ mouse has a mild or absent visual phenotype, we analyzed its retina by immunohistochemistry. We found that the *Bsg1*^−/−v2^ retina displays an increased expression of plasmalemma vesicle-associated protein (PLVAP / MECA-32), a marker used to characterize the *Nxnl2*^−/−^ mouse (**Figure 7K**) (Jaillard *et al*., 2021). The increased expression of this biomarker in vascular endothelial cells of the inner retina resembles what is observed in models of diabetic retinopathy (Sheskey et al., 2021). This observation indicates that the *Bsg1*^−/−v2^ retina is abnormal, even though there is no apparent consequence for vision under these experimental conditions.

## DISCUSSION

The RdCVF signaling of *Hydra vulgaris* is independent of BSG1, which can however be observed in the genome of the scallop. RdCVF expression is specific to ciliated photoreceptors in both scallop and amphioxus. The expression of the two products of the *NXNL* genes, RdCVF and RdCVFL, and the RdCVF receptor, BSG1, is progressively restricted to the eye, as is already achieved in the jawless vertebrate progenitors of lamprey. In the mouse, the expression of BSG1 is regulated by splicing inhibition.

### The thioredoxin activity of RdCVFL

Cellular antioxidant mechanisms arose during evolution to protect macromolecules from damage caused by reactive oxygen species (ROS) (Balsera and Buchanan, 2019). Oxidation of cysteines and methionines in proteins is of particular significance, as the thiol group can pass through reversible oxidative modifications. Most organisms share two common NADPH-dependent redox systems powered by glucose to fulfill this task: the glutathione (GSH) system and the thioredoxin system (Salinas and Comini, 2018). Thioredoxins (TXNs) are present in eukaryotes, bacteria, and archaea (Gamiz-Arco et al., 2019). The ancestral TXNs, which existed almost 4 billion years ago, display the canonical TXN fold with only minimal structural changes (Modi et al., 2018; Napolitano et al., 2019).

The catalytic mechanism of TXN involves docking to a target protein via a surface area or groove, which has been deepened during evolution to improve the specificity of substrate-enzyme interactions and for the emergence of new functions by new targeting proteins (Holmgren and Lu, 2010). Importantly, TXN family members are compartmentalized, such that mammalian TXN1 is in the cytosol and the nuclei, and TXN2 is in the mitochondria.

The RdCVFL thioredoxin fold is homologous to that of nucleoredoxin (NXN), a TXN family member with a three-tryparedoxin (TRYX)-fold, first identified in the trypanosome (Chalmel *et al*., 2007; Funato and Miki, 2007). RdCVFL and NXN have presumably a common ancestor because of the existence of plant *NXN* genes. NXN1 in plants protects cells against oxidative stress (Kneeshaw et al., 2017). In contrast, the inactivation of the *Nxn* gene in the mouse results in a cellular increase in resistance to oxidative stress (Tran et al., 2021), in agreement with the function of NRX in mammals as an oxidase rather than a reductase (Urbainsky et al., 2018).

The increased susceptibly of photoreceptors in the *Nxnl1*^−/−^ mouse shows that RdCVFL is a reductase contrary to NXN (Cronin *et al*., 2010; Elachouri *et al*., 2015; Mei *et al*., 2016), as for RdCVFLa and RdCVFLb in *Hydra vulgaris* (**Figure 1K**). *Hydra vulgaris* TXN1 is ubiquitously expressed, as is hydra RdCVFLa and RdCVFLb (Perween et al., 2021) (**Figure 1C** and **E**); but TXN and RdCVFL-targeted proteins may differ, which would explain the presence of both proteins in the hydra cytosol (Gamiz-Arco *et al*., 2019). In the vertebrate retina, RdCVFL interacts with and reduces cysteines of the cytoplasmic microtubule-binding domain of TAU (Fridlich *et al*., 2009). During evolution, TAU was derived from a microtubule-associated-protein (MAP) in a metazoan, non-vertebrate ancestor (Sundermann et al., 2016). Since microtubules are involved in elongation of the *Hydra vulgaris* body column (Takaku et al., 2011), RdCVFLa and RdCVFLb silencing may result in hydra MAP cysteine oxidation, triggering MAP release from microtubules and contraction of the hydra body (**Figure 1G** and **K**).

One of the key steps in the evolution of the *NXNL* genes is the progressive tissue-specific restriction of their expression, from a broad expression in *Hydra vulgaris* to a restricted expression in the retina of the lamprey (**Figure 6D**). The expression of *NXNL* to the distal retina of scallop indicates that the function of *NXNL* is required for only one class of photoreceptor, those with a cilium (**Figure 4F**). Microtubule alterations may destabilize photoreceptor integrity, while RdCVF2L increases the length of the rod outer segment of the *Nxnl2*^−/−^ mouse (Jaillard *et al*., 2012; Nag et al., 2020). Studies on the evolution of gene expression point to variation in both *cis* and *trans* which, under selection, can preferentially accumulate over time (Signor and Nuzhdin, 2018). In the mouse both *Nxnl1* and *Nxnl2* expression by photoreceptors is controlled though a CRX binding element, which is a *cis* element (Lambard et al., 2010). One of the shared developmental aspects of vertebrate and *Drosophila* eyes is their association with cephalization of the nervous system (Koenig and Gross, 2020). Evolutionary divergence in *cis* elements may explain why *Nxnl2* is also expressed specifically by a subset of cells of the *area postrema*, a region of the brain at the interface between blood circulation at the cerebrospinal fluid, implicated in the control of glucose supply to the brain (Jaillard *et al*., 2021).

### The trophic factor RdCVF

The main surprise in this study is the existence of intron retention of *NXNL* in *Hydra vulgaris*, a species without eyes. The first step of spliceosome assembly for all eukaryotic cells is initiation by U1 snRNP interaction with the donor site, 5’ to the intron to be spliced (Borao et al., 2021). RNA Polymerase II scans the pre-mRNA in the 5′–3′ direction as the transcript elongates, and it binds to a donor site if the interaction is sufficiently strong (Braunschweig et al., 2014). Intron retention can be simply explained by detachment of the snU1-RNP before transcription reaches the acceptor site, 3’ of the intron (Monteuuis et al., 2019). We did not detect in the retained introns of hydra *NXNLA* and *NXNLb* genes any specific sequence or secondary structure (**Figure S2B** and **C**), but the difference in the ratio RdCVFa/RdCVFLa to that of RdCVFb/RdCVFLb indicates that intron retention is certainly encoded in DNA (**Figure 2D**).

Messenger mRNAs with retained introns are generally restricted from exiting the nucleus or degraded in the cytoplasm by nonsense-mediated mRNA decay (NMD) (Grabski et al., 2021). Nevertheless, all retroviruses utilize mechanisms that allow the export and translation of mRNAs with one or more retained introns, and there are numerous examples of mRNAs with intron retention that are efficiently exported from the nucleus and used to express novel protein isoforms (Ge et al., 2016). Northern blot analysis unambiguously validates intron retention of the *NXNL1* gene in the human retina (Ait-Ali *et al*., 2015). NMD protects against the deleterious effects of truncated proteins with potential dominant-negative or gain-of-function activity (Lloyd, 2018); but the gain-of-function of RdCVFa for *Hydra vulgaris* is probably beneficial to the organism, as is demonstrated by its interaction with a putative cell surface receptor (**Figure 3J**).

Thioredoxins and RdCVF are devoid of a secretory signal and are secreted by an unconventional protein secretion pathway that has not yet been elucidated (Leveillard *et al*., 2004; Sitia and Rubartelli, 2020). Because the extracellular space is oxidized, we postulated that RdCVF may act as a pro-oxidant toward extracellular GLUT1 cysteines after RdCVF-BSG1 interaction, which triggers GLUT1 tetramerization and increases the *V_max_* of glucose uptake (Camacho et al., 2016). During evolution, when RdCVF starts to interact with a BSG1 protein that is complexed with GLUT1, this system could generate a rapid transition in accordance with the punctuated equilibrium model of evolution (Schneider, 2000). The benefit for the cell of an enhanced protection against oxidative damage is certainly a trait retained by natural selection during evolution (Leveillard and Ait-Ali, 2017). We show that this happened before the emergence of the eye (**Figure 2B**).

Cnidarians have a large range of visual organs, from eye spots to lens-containing eyes. The majority of these photoreceptive organs are found within the *Medusazoan* clade, including corals, sea anemones and jellyfish (Picciani et al., 2018). The last common *Cnidarian* ancestor, as well as ancestors of *Medusozoans* including *Hydrozoa* (which contains the clade of *Hydra vulgaris*), have no eye. Cnidocyte discharge predated eyes, perhaps facilitating the evolution of eyes in *Cnidarians* (Picciani et al., 2021). RdCVF is a module that may have been involved in cnidocyte regeneration and was recruited for photoreceptor metabolism during evolution (Swafford and Oakley, 2019). *Cnidarians* invest a large fraction of their energy in the maintenance of their cnidocyst repertoire, which has to be constantly renewed, like the outer segments of photoreceptors (Beckmann and Ozbek, 2012).

### The RdCVF receptor BSG1

The interaction of hydra RdCVFa with BSG1 from lamprey and mouse shows that the primary sequence of RdCVFa carries the information required to bind to the Ig0 domain of BSG1 even though truncated in its thioredoxin fold (Chalmel *et al*., 2007) (**Figure 2A**). All members of the BSG family have a conserved hydrophilic amino acid (E) in their transmembrane domain, which is essential for interaction with lactate transporters and probably with GLUT1 (**Figure 3A**) (Muramatsu, 2016). Both GLUT1 and MCT1 belong to the superfamily of solute carriers. The transmembrane and cytoplasmic regions of BSG are essential and sufficient for association with MCT1, so that BSG2 interacts with MCT1.

In the scallop eye, we did not detect any spliced mRNA that encodes for BSG2 (**Figure 4H**). In the amphioxus body, a BSG2 mRNA is barely detectable in the anterior part of the organism (**Figure 5G**). This finding implies that the ancestor protein encoded by the *BSG* gene is BSG1, a protein with three extracellular Ig domains that bind to RdCVF (**Figure 4I**). Glucose transporters of class I (GLUT1 and GLUT3) are specific to multicellular animals (Wilson-O’Brien et al., 2010), and glucose is the major source of energy for neurons. The scenario of the emergence of RdCVF signaling indicates an ancestral association of BSG1 with a glucose transporter rather than with a lactate transporter.

Interestingly, in the mouse eye, the production of BSG2 mRNA is controlled by splicing inhibition (**Figure 7I** and **J**), presumably involving HNRNPH2 (Grammatikakis et al., 2016; Stark et al., 2011). (**Figure S6G**). This mode of action explains the deficit of photoreceptor function of the *Bsg1*^−/−v1^ mouse through the lack of lactate transporters in the extracellular membranes of retina and RPE (Marchiq et al., 2015). Although the splicing reaction is carried out with very high precision, the *cis*-acting signals that mediate spliceosome binding show limited sequence constraint. As a result, pre-mRNAs harbor a large number of cryptic splice sites (Zarnack et al., 2013). The cell tightly controls the accessibility of such signals to the splicing machinery. The remodeling of these *cis*-acting signals allows the emergence of novel exons during evolution (Das et al., 2019).

There is little or no effect on visual function in the *Bsg1*^−/−v2^ mouse, which suggests that RdCVF can sustain cone function independent of its interaction with BSG1 (Ait-Ali *et al*., 2015) (**Figure 7E**). The expression of *Nxnl2* in the brain and the abnormal behavior of the *Nxnl2*^−/−^ mouse together indicate that the RdCVF2 cell-surface receptor is distinct from BSG1, since BSG1 expression is restricted to photoreceptors (Ochrietor et al., 2003). Consequently, there is possibly a crosstalk between the RdCVF and RdCVF2 receptors. Our comparative genomic analysis provides an argument for this crosstalk, since the BSG gene of the pig does not contain a sequence encoding for an Ig0 (**Figure S3A**). Thus even though pig BSG1 does not exist, RdCVF nevertheless protects cones in that species (Noel et al., 2021).

## Acknowledgments

Sébastien Artigaud, Marie-Christine Birling, Christopher Cabral, Carole Cossu-Leguille, Hector Escriva, Vanessa Ferracane, Stéphane Fouquet, Brigitte Galliot, Morgane Lejault, Enrico Nasi, Ousmane Niakate, Kamila Remini, Angela Tino, Yvan Wenger, the visual phenotyping and imaging platform, *Institut de la Vision*, the electron microscopy facility, *Institut de Biologie Paris-Seine*. This research was funded by Sorbonne Université, Inserm, IHU FOReSIGHT [ANR-18-IAHU-0001], Infrafrontier (K4443) and the *Fondation de France* (TL 2016 award); and (to GLF) the Great Lakes Fishery Commission, NIH Grant EY001844, an unrestricted grant from Research to Prevent Blindness USA to the UCLA Department of Ophthalmology, and NEI Core Grant EY00311 to the Stein Eye Institute.

## Author contributions

N.AA, F.B., E.C., A.M., Q.C., F.D., M.K., C.B, and X.R. conducted the experiments; J.H. conducted the experiments and helped write the paper. A.vD provided scientific expertise; A.T. provided reagents and scientific expertise; G.L.F. provided scientific expertise and helped write the paper; N.AA and T.L designed the experiments and wrote the paper.

## Declaration of interests

The authors declare no competing interests.

## Figure legends

**Figure S1. Phylogenic alignment of RdCVFL proteins.** Cysteine and serine in catalytic site (yellow and orange). In blue, conserved aspartic acid (D) or glutamic acid (E) that has a role in RdCVF binding to BSG1. The β-sheets in the secondary structures are colored in green, the α-helix in red.

**Figure S2. Phylogenic alignment of RdCVF proteins and analysis of intron retention**. (**A**) Cysteine and serine in catalytic site (yellow and orange). In blue, conserved aspartic acid (D) or glutamic acid (E) that has a role in RdCVF binding to BSG1. (**B**) Red frame, the intronic region scanned for mRNA secondary structures. The black frame shows the pyrimidine tracts. (**C**) Comparison of the sequence of *NXNLa* and *NXNLb* introns. In green, exonic nucleotides. In red, 5’ and 3’ splicing sites.

**Figure S3. Phylogenic alignment of basigin and neuroplastin proteins.** (**A**) The protein domains are underlined in yellow for the signal peptide, orange for the immunoglobulin domain, blue for the transmembrane domain, and grey for the intracellular domain. The colored amino acids in the highly conserved transmembrane domain are: in light blue, a conserved glutamic acid; in orange and green, conserved amino acids depending on the protein (alanine for BSG and proline for NPTN). (B) Structural model of RdCVFLa and RdCVFLb. (**C**) Phylogenic alignment of the sequence surrounding RdCVFa E97.

**Figure S4. The distal and proximal scallop retina and RdCVF binding to BSG1.** (**A**) Expression of Pscop2 by the distal retina by immunohistochemistry. (**B**) Expression of Gqα protein by the proximal retina. (**C**) Expression of BSG1 by scallop eyes. RT: reverse transcriptase. (**D**) Analysis of the secretion of AP-fusion proteins after transfection of HEK293 cells. Mus: mouse AP-GFP-RdCVF. Sc.: scallop AP-GFP-RdCVF. (**E**) Analysis of the expression of HA-fusion proteins after transfection of HEK293 cells.

**Figure S5. Scallop BSG1.** (**A**) Sequence of the scallop BSG1. The underlined sequences have been identified as tryptic peptides by mass spectrometry. The immunoglobulin domains are colored in red (N-terminal Ig), blue (Ig1), and green (Ig2). catalytic sites and the non-conserved E/K position 97 are indicated in red. (**B**) Analysis of the secretion of AP-fusion proteins after transfection of HEK293 cells. AP-GFP-L: lamprey BSG1 fusion protein.

**Figure S6. The *Bsg1* knock-in mice.** (**A**) representation of the *Bsg1*^−/−v1^ knock-in mouse. (**B**) genotyping of the *Bsg1*^−/−v1^ knock-in mouse. (**C**) representation of the *Bsg1*^−/−v2^ knock-in mouse. (**C**) genotyping of the *Bsg1*^−/−v2^ knock-in mouse. (**E**) Sequence of *Bsg* exon 2 with the stop codon introduced by homologous recombination. The letter A in red corresponds to the mutations. First and last position (*Bsg1*^−/−v1^ construct), second position (*Bsg1*^−/−v2^ construct). (**F**) Splicing inhibitory elements within exon 2 and the effect of the introduction of stops codons. (**G**) Expression of HNRNPF and HNRNPH2 analyzed by western blotting. RPE: retinal pigment epithelium, Ret.: retina.

**Video S1. Collection of short photoreceptors (rods) from the lamprey retina.**

**Video S2. Collection of long (cones) from the lamprey retina.**

## STAR Methods

### Animals

*Hydra vulgaris* (**Figure 1–3**) were originally obtained from Angela Tino (*Consiglio Nazionale delle Ricerche*, Naples). They were asexually cultured in Hydra medium (1 mM CaCl_2_, 0.1 mM MgCl_2_, 0.1 mM KNO_3_, 0.5 mM NaHCO_3_, 0.08 mM MgSO_4_) at pH 7. The animals were kept at 18 ± 1°C and fed three times per week with freshly hatched *Artemia salina* nauplii. Scallop eyes and tissues (**Figure 4**) were collected by the Department of Biology and Geosciences of Osaka (Japan). Mantle tissue with eyes from scallop was fixed overnight at 4°C in a solution of 4% PFA and stored at −80°C and sent with dry ice. Amphioxus (*Branchiostoma lanceolatum*) animals (**Figure 5**) were kindly provided by Hector Escriva (*Biologie intégrative des organismes marins* at the *Observatoire Océanologique at Banyuls-sur-Mer*, France). Animals were kept in a fish tank in the laboratory for 6 months at room temperature.

Lamprey experiments were performed at UCLA (**Figure 6**). Adult sea lamprey (*Petromyzon marinus*) were provided by the Hammond Bay Biological Station of the United States Geological Survey (USGS), Millersburg, MI, USA. They were captured in tributaries of Lake Huron (Ocqueoc River and Cheboygan River) in the process of their upstream spawning migration and shipped to UCLA with a permit from the Department of Fish and Wildlife of the California Natural Resource Agency. Animals were kept for no more than 6 weeks in a large tank filled with constantly circulating de-chlorinated water chilled to 4°C with an aquarium chiller (AquaEuroUSA, Gardena, CA, USA). The tank was kept in a cold room at 10°C and was maintained under 12-h light/dark room lighting. Animals were anaesthetized by immersion in 250 mg tricaine methane sulfonate, decapitated, and pithed rostrally and caudally before removal of the eyes. Care and handling of mice in these studies conformed to the rules set by the National Institutes of Health, USA, and by the Association for Research on Vision and Ophthalmology.

Care and handling of mice in these studies conformed to the rules set by the National Institutes of Health, USA, and by the Association for Research on Vision and Ophthalmology. All experiments were performed in accordance with the European Community Council Directives of September 22, 2010 (2010/63/UE). *Bsg1*^−/−v1^ and *Bsg1*^−/−v2^ knock-in mice were generated by Christine Birling (*Institut Clinique de la Souris*, Strasbourg, France) by homologous recombination (**Figures 7** and **S6**) on a pure C57BL/6J background. Mice were genotyped using multiplex PCR primers (5’-3’) TGTAGAACCAGATGCTGCTCTGCCCTA, GGCTCTGCAAAGTCTGGCTGGAA, GAAGTTATACTAGAGCGGCCGTTCAC, GTGGCTGT CCTTCCTTGTAGTAACGGGTA, CAGCTCATTCCTCCCACTCATGATC, TGCTAAAGCGCA TGCTCCAGACTGC and GGAGCAGCTGTCGTTTGGAGCAT. The mouse lines were maintained at the animal facility Charles Foix (UMS28) under standard conditions with access *ad libitum* to food and water and with a 12-h light/dark cycle. The animals under experimentation were transferred to the animal facility of the Institut de la Vision for 5 years under the agreement obtained April 26^th^ 2016 between the *Direction Départementale De La Protection Des Populations De Paris* (B-75-12-02) and principal investigator (T.L.), certificate (N°A-75-1863; OGM n°5080 CA-II). Mice were housed with access *ad libitum* to food and water with a 12-h light/dark cycle of 20-50 lx.

### Phylogenetic analysis

The protein sequences (**Figure S1**) were retrieved from the appropriate public databases (**Table S1**). The hydra sequences were extracted from the [NCBI *Hydra vulgaris* Annotation Release 102](https://www.ncbi.nlm.nih.gov/genome/annotation_euk/Hydra_vulgaris/102/) (XP_004206112.1, XP_002165519.2). The amphioxus reference protein sequences were extracted from [Amphiencode](http://amphiencode.github.io/Data/), then a BLAST was performed with Hydra orthologs to select the candidate sequences (BL21340_evm0, BL21335_evm0, BL10540_evm1, BL07298_evm1). The same approach was used for the ctenophore sequences, extracted and blasted from [the MGP portal](https://kona.nhgri.nih.gov/mnemiopsis/) (ML104337a, ML055026a, ML093031a, ML104336a). The scallop sequences were extracted from the [*Mizuhopecten yessoensis* Annotation Release 100](https://www.ncbi.nlm.nih.gov/genome/annotation_euk/Mizuhopecten_yessoensis/100) (XP_021363738.1). [Uniprot](https://www.uniprot.org/) was used to extract the lamprey (S4RDF3), human (Q96CM4, Q5VZ03), macaque (A0A2K5WBF2, A0A2K5VRU3), mouse (Q8VC33, Q9D531), rat (F1LP37, D4A212), xenopus(Q68EV9, Q6GQZ1), zebrafish (A5PMF7, Q5PRB4) and pig (A0A287AW42, A0A286ZMV1) sequences. The 3D structures were then modeled with SWISS-Model (https://swissmodel.expasy.org/interactive) and downloaded as PDB files. The models were loaded in Chimera (https://www.cgl.ucsf.edu/chimera/), then aligned with the MatchMaker tool. The structural alignment was inferred with the Match≥Align tool. The sequences between the aligned structures were re-aligned with Clustal Omega (https://www.ebi.ac.uk/Tools/msa/clustalo/) for better readability. Jalview (https://www.jalview.org) was used to display and colorize the final alignment, and the phylogenetic tree was calculated with the standard protein matrix PAM250. Species used in the alignment were as follows, amphioxus: *Branchiostoma lancealatum*; ctenophore: *Mnemiopsis leidyi*; human: *Homo sapiens*; hydra: *Hydra vulgaris*; lamprey: *Petromyzon marinus*; macaque: *Macaca fascicularis*; mouse: *Mus musculus*; pig: *Sus scrofa*; rat: *Rattus norvegicus*; scallop: *Mizuhopecten yessoensis*; xenopus: *Xenopus laevis*; zebrafish: *Danio rerio*. Note that the lamprey sequence does not start with a methionine, which we believe is due to the loss of part of its genome in adult animals (Smith et al., 2009).

The protein sequences (**Figure S2A**) were retrieved from the appropriate public databases. The hydra sequences were extracted from the NCBI *Hydra vulgaris* Annotation Release 102 (XP_004206112.1, XP_002165519.2). The amphioxus reference protein sequences were extracted from Amphiencode, then a BLAST was performed with the Hydra orthologs to select the candidate sequences (BL21340_evm0, BL21335_evm0, BL10540_evm1, BL07298_evm1). The scallop sequences were extracted from the *Mizuhopecten yessoensis* Annotation Release 100 (XP_021363738.1). Uniprot was used to extract the lamprey (S4RDF3), human (Q96CM4, Q5VZ03), macaque (A0A2K5WBF2, A0A2K5VRU3), mouse (Q8VC33, Q9D531) and rat (F1LP37, D4A212) sequences. The short forms of these sequences were experimentally validated with RT-PCR then aligned with Clustal Omega (https://www.ebi.ac.uk/Tools/msa/clustalo/). Jalview (https://www.jalview.org) was used to display and colorize the final alignment, and the phylogenetic tree was calculated with the standard protein matrix PAM250. The 5’ and 3’ ends of the introns from the 2 *NXNL* sequences from *Hydra vulgaris* were displayed and colorized in Jalview. The sequence compositions for the pyrimidine tract were compared with each other. The secondary structure of the 5’ ends was modeled with the RNAfold web server (http://rna.tbi.univie.ac.at/cgi-bin/RNAWebSuite/RNAfold.cgi). The sequences are colorized according to the probability of being in that state (paired/unpaired), red being the highest probability. The retained introns 4 do not have sequence conservation and do not significantly differ from the others in term of secondary structure 3’ to the 5’ splicing site and to the content of pyrimidine or thymidine (%TC: 62.4% ±6) in the pyrimidine tracts 5’ to the 3’ splicing sites. Note that in a small fraction of donor sites, GT is replaced by GC (Churbanov et al., 2008), as here for *NXNLa* intron 3.

The protein sequences (**Figure S3**) were retrieved from the appropriate public databases. We selected the longest isoform available containing the Ig0 domain, when available. The hydra sequence was extracted from the NCBI *Hydra vulgaris* Annotation Release 102 (XP_002161571.3). The scallop sequence was extracted from the *Mizuhopecten yessoensis* Annotation Release 100 (XP_021372255.1). The amphioxus reference protein sequences were extracted from Amphiencode, then a BLAST was performed with several orthologs to select the candidate sequence (BL11268_cuf1). Uniprot was used to extract the lamprey (S4RAQ8), human (P35613, Q9Y639), macaque (A0A2K5WZU2, A0A2K5W4W3), mouse (P18572, P97300), rat (P26453, P97546), xenopus (A1L3J1, A0JMY2), zebrafish (Q1LVH3, Q6NZV3) and pig (A0A286ZS77, F1SIC5) sequences. The domains were defined with Interpro. The sequences from each domain were then aligned together with Clustal Omega (https://www.ebi.ac.uk/Tools/msa/clustalo/), and all sequences were pasted back together. Jalview (https://www.jalview.org) was used to display and colorize the final alignment. The pig BSG and the xenopus NPTN do not possess the first Ig domain (Ig0), preventing these species from being integrated into the tree at their proper position.

### Hydra morphology scored after copper exposition

Four groups of 10 hydra polyps (**Figure 1G** and **H**) were collected in plastic multiwells and allowed to equilibrate at room temperature in 1 ml of Hydra medium (HM) at pH 7.0. Increasing concentrations of copper were added, and copper toxicity was determined by analyzing hydra polyp morphological changes according to score values (ranging from 1 to 10) described previously (Wilby et al., 1990).

### Hydra *in vivo* RNA interference and copper exposition

Four groups of 10 hydra polyps (**Figure 1I-K**) were collected in plastic multiwells and allowed to equilibrate at room temperature in 1 ml of hydra medium at pH 4.0. Hydra polyps were incubated under a normal feeding regime for 48 h with 70 nM siRNA targeting hydra RdCVFLa (5’-GCCAGGAAGATATGATAAA-3’) labeled 3’ with Alexa fluor 488, and RdCVFLb (5’-GACTTTGTTCAAAGCTGAT-3’) labeled 3’ with Alexa fluor 647. The non-targeting siRNA (5’-ATTGATCAGAACAGAGAAT-3’) labeled 3’ with Alexa fluor 488 was used as a control. siRNAs were designed with the RNAi ThermoFisher service and purchased by QIAGEN. Hydra polyps were washed with ultrapure water 4 times and incubated in HM at pH 7.0 with 4.5 nM copper. Hydra polyp morphology were scored as previously described.

### RNA Purification, reverse transcription-PCR and qRT-PCR analysis

Depending on the tissue mass, total RNA was purified with either an RNeasy Mini or a Nano kit (Qiagen) (**Figures 1–7**). RNA was reverse transcribed with the Superscript© II Reverse Transcriptase Kit (Life-Technologies) and subjected to RT analysis. PCR was performed with cDNA. Cycling conditions were as follow: 94°C for 5 min followed by 35 cycles of amplification (94°C denaturation for 30 s, annealing for 30 s at a temperature range of 52–58°C, and 72°C elongation for 1 min), with a final incubation at 72°C for 3 min. Samples were analyzed by quantitative PCR with power SYBR green PCR master mix (Life-Technologies) in a 7500 standard real-time PCR system (Applied Biosystems) with specific primers (**Table S2**).

### BaseScope and RNAScope *in situ* hybridization

For the BaseScope™ Assay (**Figure 1C**, **2EF** and **4F**), hydra polyps were fixed in 4% paraformaldehyde (PFA) for 24 h at room temperature (RT). They were washed in PBST (1x PBS, 0.1% Tween 20) at RT for 10 min followed by dehydration in a graded series of methanol concentrations (25%, 50%, 75% and 100%), with gentle agitation at room temperature for 10 min each. Hydra polyps could be stored in 100% methanol (MeOH) up to 1 month at −20°C before continuing if necessary; storage appeared not to affect the assay. Hydra polyps were rehydrated in a reverse MeOH/PBT series and washed for 10 min each in PBST+ 1% bovine serum albumin (BSA). Hydra polyps were incubated in 1 ml of fresh 1x Target Retrieval at 100°C for 15 min and incubated in PBST for 1 min, then in 100% MeOH for 1 min. Hydra polyps were then washed in PBST+ 1% BSA, treated with Protease III solution (ACD 322337) for 30 min at 40°C, and stored in 300 µl of Probe Diluent (ACD 300041). Probe solutions *Hydra RdCVFLa* and *b* (a 20ZZ); *Hydra Opsin2* (a 13ZZ); *Hydra TXN2* (a 10ZZ), and negative control (ACD 310043) were pre-warmed at 40°C for 10 min, gently mixed, and briefly centrifuged. Up to three different probes/channels were used (C1–C3). Hydra polyps were covered by 4 drops of probe solution and incubated overnight at 40°C. For amplification and label probe hybridization, hydra polyps were washed with 1x wash buffer (WB) for 10 min at room temperature and incubated with Amp1 pre-amplifier hybridization solution for 30 min at 40°C. Hydra polyps were washed with 1x WB for 10 min at room temperature and incubated with Amp2 pre-amplifier hybridization solution for 15 min at 40°C. Hydra polyps were washed with 1x WB for 10 min at room temperature and incubated with Amp3 pre-amplifier hybridization solution for 30 min at 40°C. Hydra polyps were washed with WB for 10 min at room temperature and incubated with Amp4 pre-amplifier hybridization solution for 15 min at 40°C. Then they were washed with WB for 10 min at room temperature and incubated with Amp5 pre-amplifier hybridization solution for 30 min at 40°C. Next they were washed with WB for 10 min at room temperature and incubated with Amp6 pre-amplifier hybridization solution for 15 min at 40°C. Finally, hydra polyps were incubated with 120 µl of RED solution (1:60 ratio of Fast RED-B to Fast RED-A) for 10 min at room temperature. Hydra polyps were mounted with mowiol (Sigma) and subjected to nanozoomer analysis. For the RNAscope® Assay, the probes used are hydra RdCVFa/b, targeted intron number 4 (a 20 ZZ); *Hydra Opsin2* (a 13 ZZ); *Hydra TXN* (a 10 ZZ); and negative control (ACD 310043). The sequences of the riboprobes are listed below (**Table S3**)

### MS/MS Analysis

Proteomic analysis was performed as previously described (Ait-Ali *et al*., 2015). François Delalande: PRIDE.

### Field emission scanning electron microscopy

Hydra polyps (**Figure 1F** and **2G**) were fixed in 4 % glutaraldehyde and 0.1 M phosphate buffer (0.1 M Na_2_HPO_4_, 0.1M NaH_2_PO_4_ at pH 7.4) for 30 min at 4°C and washed 4 times in 0.2 M phosphate buffer. The fixed hydra polyps were dehydrated in increasing concentrations of ethanol (50%, 70%, 95%, and 100%) and freeze-dried in hexamethyldysilazane. After sputter-coating with platinum, hydra polyps were observed by field-emission scanning electron microscopy (FESEM, Cambridge CamScan CS-300, England).

### Alkaline phosphatase fusion protein binding assays

The alkaline phosphatase fusion-protein binding assay (**Figure 3G-H**, **4J** and **6J**) is described in Aït-Ali et al. (2015) and was originally adapted from (Flanagan and Cheng, 2000). Sequences of hydra RdCVFa (XP_004206112.1, 104 aa), hydra RdCVFb (XP_002165519.2, 104 aa) scallop RdCVF (XP_021363738.1, 108 aa), lamprey RdCVF (S4RDF3, 110 aa) and mouse RdCVF (Q8VC33, 109 aa), preceded by a GFP sequence without initiating and stop codons, were introduced into the pAPtag-5, C-terminal to the coding sequence of alkaline phosphatase (AP, 535 aa) and missing its stop codon between the XhoI and the XbaI sites. Hydra NCAM1 (XP_002161571.3) (410 aa), scallop BSG1 (XP_002161571.3) (405 aa), Lamprey BSG1 (S4RAQ8) (367 aa) and mouse BSG1(P18572) (398 aa) tagged by HA were introduced into pcDNA3 (ThermoFisher) between BamH1 and Xho1 sites. All the plasmids were generated by Genecust. The fusion proteins were produced by transient transfection in HEK293 MSR cells (Invitrogen) with 24 µg of recombinant plasmids by using lipofectamine 2,000 (Invitrogen). The cells were grown for 48 h in Opti-MEM medium (Invitrogen). Heparin (11 nM) was added to the media for 30 min, and the conditioned medium was collected and concentrated 10-fold by ultrafiltration on filters with a cut-off threshold of 5 kDa (Amicon Ultra-15). The conditioned media were analyzed enzymatically for AP activity with para-nitrophenylphosphate as a substrate and by western blotting with anti-AP, anti GFP, and anti-HA antibodies. COS-1 cells were transfected with 1.6 µg of the corresponding receptor plasmids by using lipofectamine 2,000 (Invitrogen) in Opti-MEM medium (Invitrogen). Five hours after transfection, the medium was replaced with DMEM 10% fetal calf serum (FCS), and the cells were grown for 48 h. The cells were then washed with Hanks’ balanced salt solution (HBSS), which was free of Ca^2+^ and Mg^2+^ and contained 0.5 mg/ml BSA and 20 mM HEPES (pH 7.0) (binding buffer). They were incubated for 90 min at room temperature with a 1/20 dilution of the conditioned media containing the fusion proteins in 20 mM HEPES (pH 7.0). The cells were washed 5 times for 10 min each with binding buffer and fixed for 30 s with 60% acetone, 3% PFA in 20 mM HEPES (pH 7.0). The cells were rinsed for 10 min with HBSS 150 mM NaCl, 20 mM HEPES (pH 7.0) and warmed for 20 min at 65°C. The cells were washed in HBSS containing 100 mM Tris pH 9.5, 100 mM NaCl and 5 mM MgCl_2_. The colorimetric reaction was performed in binding buffer with 10 mM L-homoarginine, 0.17 mg/ml 5-bromo-4-chloro-3-indolyl phosphate (BCIP) and 0.33 mg/ml nitro blue tetrazolium (NBT) in the dark. The reactions were stopped in PBS containing 500 mM EDTA, and the cells were stored at 4°C.

### Lamprey (*Petromyzon marinus*) retina dissection and single-photoreceptor cell preparation

Eyes were enucleated under dim red light (**Figure 6E**). The anterior portion of the eye was cut and the lens and cornea were removed with infrared image converters. The retina was isolated from the eyecup; the retinal pigment epithelium was removed with fine tweezers, and the retina was chopped into small pieces with a razor blade. The retinas were exposed for 3 min to 0.5 mg/ml collagenase and 0.33 mg/ml hyaluronidase to prevent clogging of pipettes by vitreous humor and extracellular matrix. Single cones previously referred to as ‘long’ photoreceptors and single shorter rods (Dickson and Graves, 1979) were selected and picked up with appropriate pipettes. The resistance of useable pipettes was about 3-4 MΩ for rods and 2.5-3.0 MΩ for cones.

### Lamprey single-cell RT-PCR

cDNA of each single cell was amplified by incubating in 15 μl of RT mix (1x rRNasin® RNase inhibitor 0.75 μl (N2511 Promega); 1 x Buffer; 10 mM each dNTP mix (4303441, ThermoFisher); 0.1 M DTT 3 μl; 30 μg Random Primers (C1181, Promega), and Superscript II (18064014, ThermoFisher) 0.3). Nested-PCR was performed with cDNA. Cycling conditions were as follow: 94°C for 5 min, followed by 35 cycles of amplification (94°C denaturation for 30 s, annealing for 30 s at a temperature range of 52-58°C, and 72°C elongation for 1 min) with a final incubation at 72°C for 3 min. Primer sequences are provided in **Table S2**.

### Western blotting

Deglycosylation of 50 µg of mouse retinal extract or RPE (**Figure 7G** and **H**) was performed according to Aït-Ali et al. (2015), with a combination of the following enzymes: O-glycosidase, PNGase F, neuraminidase, β1-4 galactosidase and β-N-acetylglucosaminidase (New England BioLabs # P6039S) after the manufacturer’s instructions. The untreated extracts were incubated similarly without the enzymes. The extracts were analyzed by western blotting by using rat monoclonal anti-basigin (anti-CD147 Abcam ab34016, 1/350). To check the expression of the fusion proteins, we sonicated transfected COS-1 cells twice for 10 s on ice in 50 mM Tris-HCl (pH 7.5), 1 mM EDTA, 1 mM DTT, 1% Triton X-100, 1 mM phenylmethylsulfonyl fluoride (PMSF), and 0.14 mM tosyl-L-lysine chloromethyl ketone hydrochloride (TLCK), in the presence of a cocktail of proteinase inhibitors (P2714, Sigma, St Louis, MI, USA). After 30 min on ice, the extracts were centrifuged for 5 min at 12,000 rpm at 4°C. We resolved 40 µg of whole-cell extract on a NuPAGE® Novex 4-12% Bis-Tris (NP0322BOX, Invitrogen) polyacrylamide gel in NuPAGE® MES SDS Running Buffer (NP0002, Invitrogen) at 180 V and rapidly transferred onto nitrocellulose membrane, 0.2 µm (GE Healthcare). After saturation, the membranes were incubated with anti-BSG antibody (Abcam, Ab34016) (**Figure 7GH**).

After saturation, the membranes were incubated (**Figure 3E, F**, **S4D and S5B**) with rabbit polyclonal AP antibody, provided with the AP Western Blot Kit (Q310, GenHunter), anti-GFP antibody (Roche Cat# 11814460001, RRID: AB_390913), anti-HA antibody (Covance, catalogue number MMS-101R-0500; RRID: AB_2314672). For western blotting analysis of (**Figure S6G**) the membranes were incubated with rabbit polyclonal anti HNRNPF antibody (Abcam Cat# ab50982, RRID: AB_880477) and rabbit polyclonal anti HNRNPH2 antibody (Abcam Cat# ab84744, RRID: AB_1925071).

### Far-western blotting

Far-western blotting (**Figure 3I** and **J**) was performed as previously described (Ait-Ali *et al*., 2015). The *E. coli* codon optimized open reading frame of hydra-RdCVFa (104 residues) and hydra-RdCVFb (104 residues) without their respective methionine initiation codons were cloned in a pET41TEV plasmid vector with a N-terminus His-GFP-TEV fusion tag and a 6 histidine (His) tag 6His-GFP-SUMO-hydraRdCVFa or b. One liter of *E. coli* culture was used to purify 25 mg of protein on Ni-charged immobilized metal ion affinity chromatography (IMAC) in 50 mM Tris pH7.5, 250 mM NaCl, 20 mM imidazole, 2 M urea, and 0.1 mM PMSF followed by gel filtration. Membrane and soluble fractions from *Hydra vulgaris* and COS-1 cells were prepared with FOCUS™ membrane proteins (G-Biosciences); total fraction (T) was prepared with the total protein extraction kit (G-Biosciences).

### Immunohistochemistry

After fixation, scallop eyes (**Figure 4D, E, S4A, S4B** and **Table S4**) were isolated from mantle tissue and prepared immediately for cryosectioning by overnight incubation at 4°C in 10%, 20%, and 30% sucrose solutions (in PBS). With the cryostat (Thermo Scientific MicromHM505-N) set to −20°C, scallop eyes were sectioned at 8 μm, and sections were collected onto slides (superfrost, Thermoscientific) and stored at −20°C. First sections were permeabilized at room temperature in PBS with 0.3% Triton X-100, for 10 min, three times. Sections were blocked in PBS-TX with 10% normal goat serum at room temperature for 1 h (blocking buffer). Sections were incubated with primary rabbit polyclonal Anti-Gq (Merck, 371751) antibodies against Gqα (1/200)and pScop2 (1/500), kindly provided by Enrico Nasi (*Universidad Nacional de Colombia*, Bogota, Colombia) (Arenas *et al*., 2018), and were diluted in blocking buffer. Slides were then covered with strips of parafilm and stored at 4°C overnight. After three washes with PBS, the slides were incubated with goat anti-rabbit Ig antibody fluo-488 and goat anti-rabbit Ig antibody fluo-594 1 h at room temperature. After 3 additional washes in PBS, the slides were incubated 10 min with 36 mM DAPI. Finally, sections were washed three times with PBS before mounting with Fluoromount-G (SouthernBiotech) and coverslips were sealed with nail polish. Images were obtained with a confocal microscope (Olympus FV1000). Stacks of consecutive images taken at 500 nm intervals were acquired sequentially with lasers: diode 559 nm, argon 488 nm and diode 405 nm. Z projections of serial sections were reconstructed in the FV-10-ASW-4.2 software of the confocal microscope (**Figure 7K**). Immunostaining of 12-μm retinal sections of *Bsg1*^+/+^, *Bsg1*^−/−v1^and *Bsg1*^−/−v2^ mice at PN40 (**Figure 7L**) were prepared by incubation with Rat Mouse MECA-32 antibody (DSHB Cat# MECA-32, RRID: AB_531797) and DAPI. Images were viewed on a Zeiss Axiovert 100 inverted microscope.

### Scotopic and photopic electroretinography

Mice were anesthetized intraperitoneally with a mixture of ketamine (80 mg/kg) and xylazine (10 mg/kg) diluted in saline (**Figure 7A-C** and **E**). Their pupils were dilated by topical application of 0.5% mydriaticum. Body temperature was maintained at 37°C with a circulating hot water heating pad. The electrical signal was recorded with a pair of electrodes fixed to our platform and adapted for use on mice. A gold loop electrode was placed at the center of the corneal surface and maintained in position with ocrygel (TVM) to ensure good electrical contact. Two stainless steel reference electrodes were inserted subcutaneously in each cheek of the mouse to normalize signal output, and the ground electrode was inserted subcutaneously in the back of the mouse. Recordings were made from both eyes. The light stimulus was provided by a 150 W xenon lamp in a Ganzfeld stimulator (Multiliner vision, Jaeger Toennies, Germany). Responses were amplified and filtered (1 Hz-low and 300 Hz-high cut off filters) with a 1 channel DC-/AC-amplifier. To isolate cone responses under photopic conditions, a 5-min saturating light at 10 cd.s/m^2^ was used to desensitize the rods. The cone photopic electroretinograms (ERGs) we show are the averages of 10 responses from 10 consecutive flashes at an intensity of 10 cd.s/m^2^. Photopic *b*-wave amplitudes were measured from the base-line to the peak. Photopic *b*-wave amplitudes were measured from the base-line to the peak of the response. For scotopic recordings, single flash responses obtained at light intensities of 3×10^−3^, 3×10^−2^, 3×10^−1^, 3 and 10 cd.s/m^2^ using a sampling frequency of 5 kHz, a flash duration of 4 ms, and a frequency of stimulus of 0.5 Hz. Data were recorded from 50 ms before stimulus onset to 450 ms post-stimulus. The amplitudes of *a-* and *b-*waves were measured from the baseline to the peak of the response.

### Spectral-domain optical coherence tomography

Mice were anesthetized and pupils were dilated (**Figure 7D**). Eye dehydration was prevented by regular instillation of sodium chloride drops. Spectral-domain optical coherence tomography (SD-OCT) images were recorded on the right eye with a spectral domain ophthalmic imaging system (Bioptigen 840 nm HHP; Bioptigen; North Carolina USA). We performed rectangular scans consisting of a 2 mm by 2 mm perimeter with 1000 A-scans per B-scan with a total of 100 B-scans. Scans were obtained while centered on the optic nerve. SD-OCT scans were analyzed with Bioptigen Diver Software 3.4.4. and layers were measured using automatic segmentation. Each analysis was reviewed for correct layering. Average thicknesses for the ONL from each quadrant were used for analysis. Outer nuclear layer (ONL) thickness was measured every 100 µm, ventral-dorsal and temporal-nasal, beginning 100 μm from the center (optic nerve).

### Cone density measurement

Measurements of cone density (**Figure 7F**) was performed with the platform e-conome after staining the cones with peanut agglutinin (PNA) (Clerin *et al*., 2011). Dissection was followed by the labeling of retinas with PNA-λ594 at 1/40 overnight. Flat-mounting was made to proceed to automatic acquisition and counting with the platform e-conome.

---

**Table S1:** Accession number of proteins

| Figure | Acronym | Source | Identifier |
| --- | --- | --- | --- |
| S1 | sponge_RdCVFL | https://www.ncbi.nlm.nih.gov/geo/ | c97660 |
|  | ctenophore_RdCVFLa | https://research.nhgri.nih.gov/mnemiopsis/ | ML104337a |
|  | ctenophore_RdCVFLb | https://research.nhgri.nih.gov/mnemiopsis/ | ML055026a |
|  | ctenophore_RdCVFLc | https://research.nhgri.nih.gov/mnemiopsis/ | ML093031a |
|  | ctenophore_RdCVFLd | https://research.nhgri.nih.gov/mnemiopsis/ | ML104336a |
|  | hydra_RdCVFLa | https://www.ncbi.nlm.nih.gov/protein | XP_004206112.1 |
|  | hydra_RdCVFLb | https://www.ncbi.nlm.nih.gov/protein | XP_002165519.2 |
|  | scallop_RdCVFL | https://www.ncbi.nlm.nih.gov/protein | XP_021363738.1 |
|  | amphioxus_RdCVFLa | https://amphiencode.github.io/ | BL21340_evm0 |
|  | amphioxus_RdCVFLb | https://amphiencode.github.io/ | BL21335_evm0 |
|  | amphioxus_RdCVFLc | https://amphiencode.github.io/ | BL10540_evm1 |
|  | amphioxus_RdCVFLd | https://amphiencode.github.io/ | BL07298_evm1 |
|  | lamprey_RdCVFL | https://www.uniprot.org/uniprot/ | S4RDF3 |
|  | zebrafish_RdCVFL | https://www.uniprot.org/uniprot/ | A5PMF7 |
|  | zebrafish_RdCVF2L | https://www.uniprot.org/uniprot/ | Q5PRB4 |
|  | xenopus_RdCVFL | https://www.uniprot.org/uniprot/ | Q68EV9 |
|  | xenopus_RdCVF2L | https://www.uniprot.org/uniprot/ | Q6GQZ1 |
|  | mouse_RdCVFL | https://www.uniprot.org/uniprot/ | Q8VC33 |
|  | mouse_RdCVF2L | https://www.uniprot.org/uniprot/ | Q9D531 |
|  | rat_RdCVFL | https://www.uniprot.org/uniprot/ | F1LP37 |
|  | rat_RdCVF2L | https://www.uniprot.org/uniprot/ | D4A212 |
|  | pig_RdCVFL | https://www.uniprot.org/uniprot/ | A0A287AW42 |
|  | pig_RdCVF2L | https://www.uniprot.org/uniprot/ | A0A286ZMV1 |
|  | macaque_RdCVFL | https://www.uniprot.org/uniprot/ | A0A2K5WBF2 |
|  | macaque_RdCVF2L | https://www.uniprot.org/uniprot/ | A0A2K5VRU3 |
|  | human_RdCVFL | https://www.uniprot.org/uniprot/ | Q96CM4 |
|  | human_RdCVF2L | https://www.uniprot.org/uniprot/ | Q5VZ03 |
| S2 | hydra_RdCVFa | https://www.ncbi.nlm.nih.gov/protein |  |
|  | hydra_RdCVFb | https://www.ncbi.nlm.nih.gov/protein |  |
|  | scallop_RdCVF | https://www.ncbi.nlm.nih.gov/protein |  |
|  | amphioxus_RdCVFa | https://www.ncbi.nlm.nih.gov/protein |  |
|  | lamprey_RdCVFL | https://www.ncbi.nlm.nih.gov/protein |  |
|  | mouse_RdCVF | https://www.ncbi.nlm.nih.gov/protein |  |
|  | mouse_RdCVF2 | https://www.uniprot.org/uniprot/ | Q9D531-4 |
|  | rat_RdCVF | https://www.ncbi.nlm.nih.gov/protein |  |
|  | rat_RdCVF2 | https://www.ncbi.nlm.nih.gov/protein |  |
|  | macaque_RdCVF | https://www.ncbi.nlm.nih.gov/protein |  |
|  | macaque_RdCVF2 | https://www.ncbi.nlm.nih.gov/protein |  |
|  | human_RdCVF | https://www.ncbi.nlm.nih.gov/protein |  |
|  | human_RdCVF2 | https://www.ncbi.nlm.nih.gov/protein |  |
| S3A | hydra_NCAM1 | https://www.ncbi.nlm.nih.gov/protein | XP_002161571.3 |
|  | scallop_BSG | https://www.ncbi.nlm.nih.gov/protein | XP_021372255.1 |
|  | amphioxus_BSG | https://amphiencode.github.io/ | BL11268_cuf1 |
|  | lamprey_BSG | https://www.uniprot.org/uniprot/ | S4RAQ8 |
|  | zebrafish_BSG | https://www.uniprot.org/uniprot/ | Q1LVH3 |
|  | xenopus_BSG | https://www.uniprot.org/uniprot/ | A1L3J1 |
|  | mouse_BSG | https://www.uniprot.org/uniprot/ | P18572 |
|  | rat_BSG | https://www.uniprot.org/uniprot/ | P26453 |
|  | macaque_BSG | https://www.uniprot.org/uniprot/ | A0A2K5WZU2 |
|  | human_BSG | https://www.uniprot.org/uniprot/ | P35613 |
|  | zebrafish_NPTN | https://www.uniprot.org/uniprot/ | Q6NZV3 |
|  | mouse_NPTN | https://www.uniprot.org/uniprot/ | P97300 |
|  | rat_NPTN | https://www.uniprot.org/uniprot/ | P97546 |
|  | pig_NPTN | https://www.uniprot.org/uniprot/ | F1SIC5 |
|  | macaque_NPTN | https://www.uniprot.org/uniprot/ | A0A2K5W4W3 |
|  | human_NPTN | https://www.uniprot.org/uniprot/ | Q9Y639 |
|  | pig_BSG | https://www.uniprot.org/uniprot/ | A0A286ZS77 |
|  | xenopus_NPTN | https://www.uniprot.org/uniprot/ | A0JMY2 |

**Table S2:**
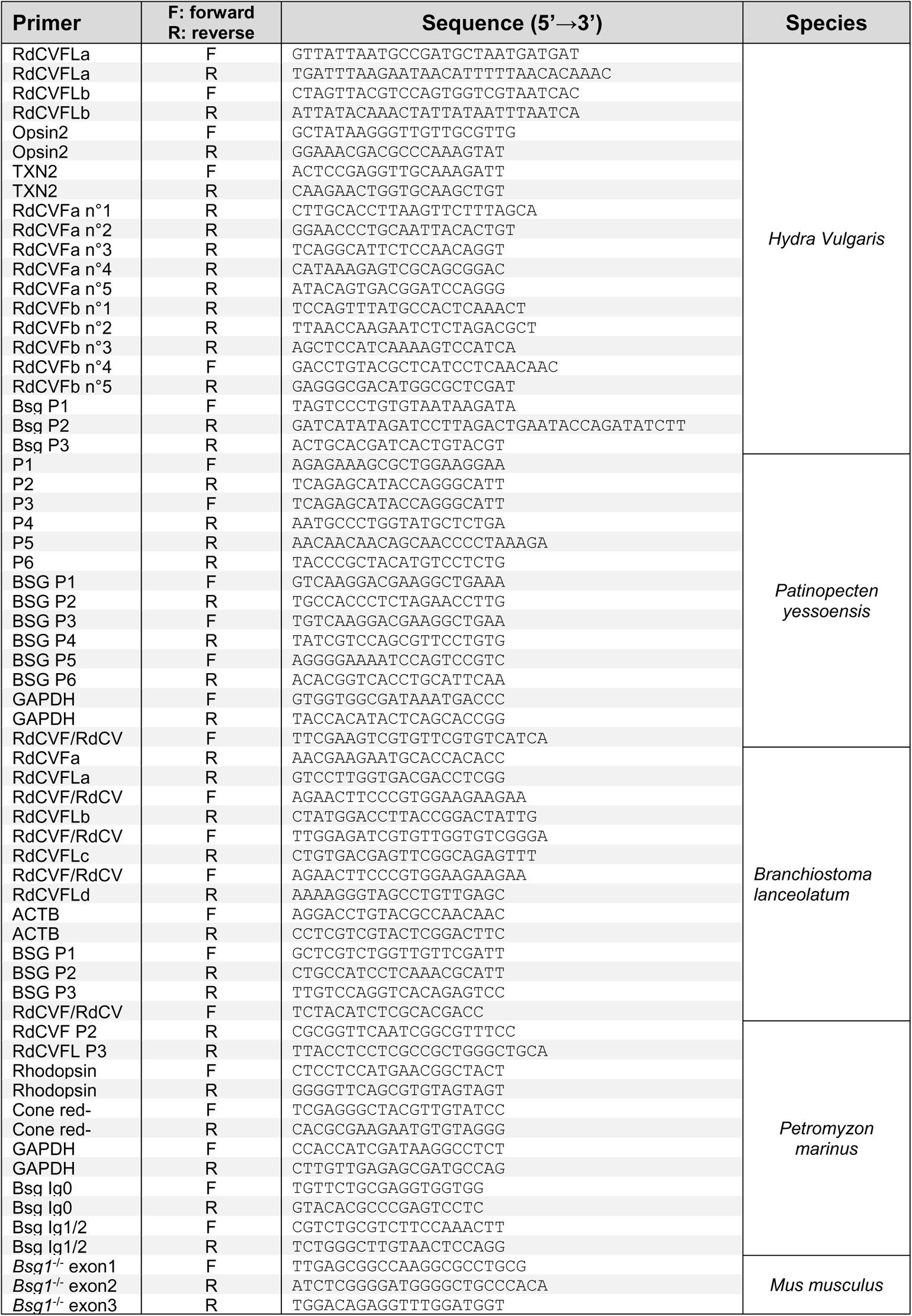
PCR primer sequences

**Table S3:**
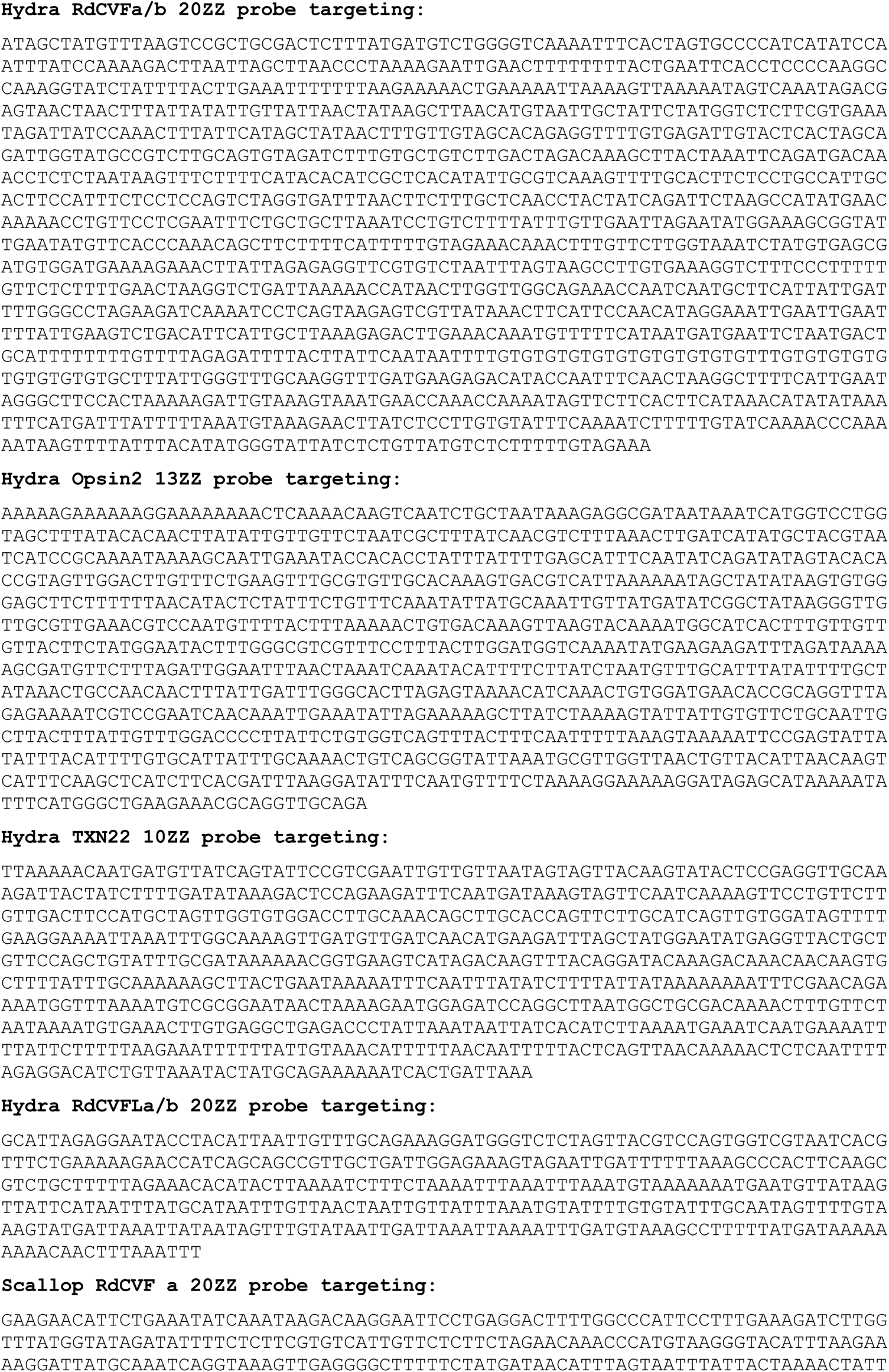
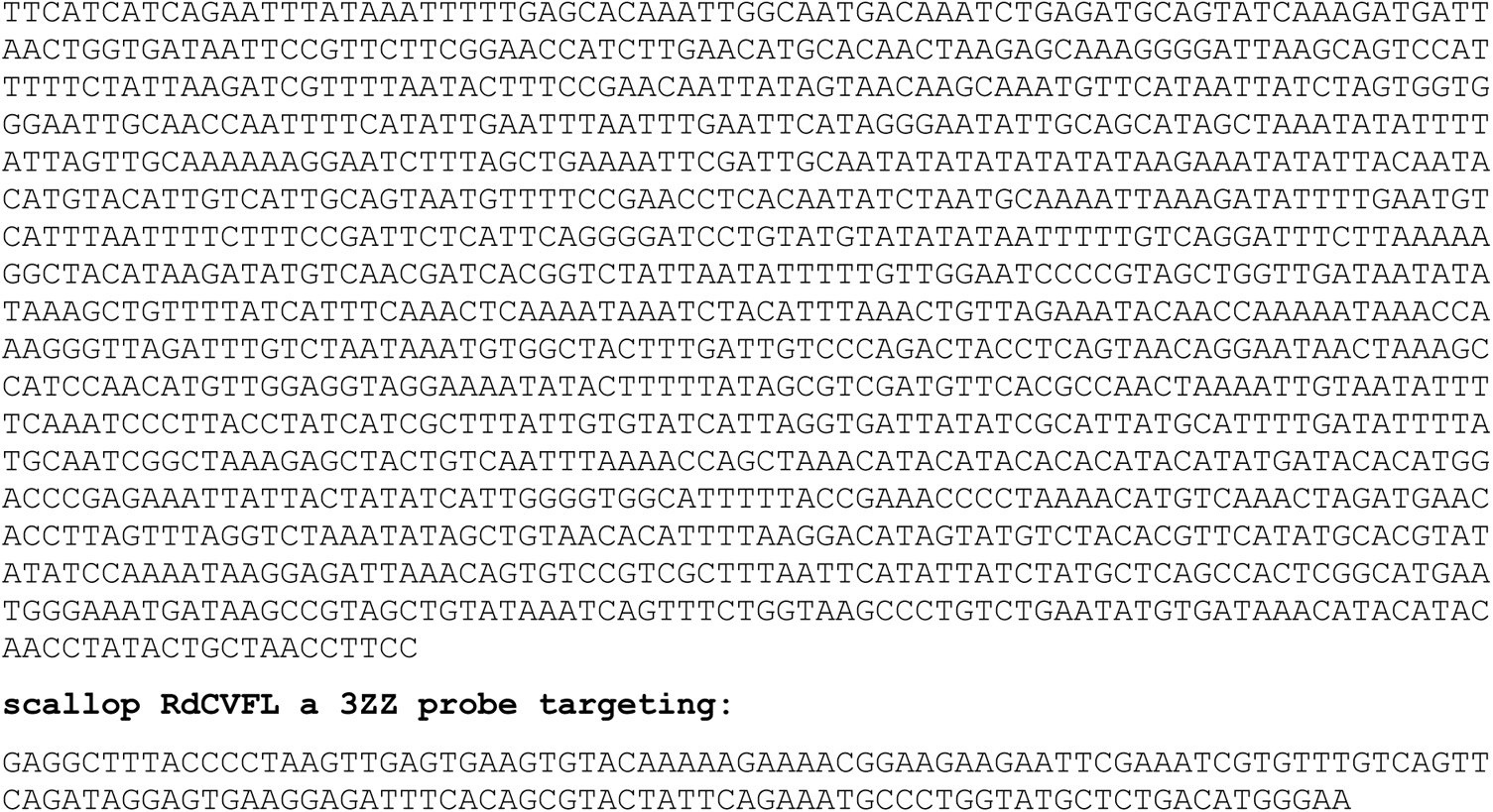
Riboprobe sequences

**Table S4:** Antibodies

| Target | Company | Catalogue number | Dilution | RRID# |
| --- | --- | --- | --- | --- |
| GFP | Roche | MMS-101R-0500 | 1/1000 | AB_2314672 |
| AP | GenHunter | Q310 | 1/500 | - |
| HA | Covance | MMS-101R-05 | 1/1000 | AB_2314672 |
| Gqα | Merck | 371751 | 1/200 | - |
| pScop2 | - | Enrico Nasi | 1/500 | - |
| BSG | Abcam | Ab34016 | 1/350 | AB_726136 |
| HNRNPF | Abcam | ab50982 | 1/1000 | AB_880477 |
| HNRNPH2 | Abcam | ab84744 | 1/1000 | AB_1925071 |
| MECA-32 | BD Biosciences | 553849 | 1/500 | AB_395086 |

## Notes

### Competing Interest Statement

The authors have declared no competing interest.

