## Supplementary figures for "The Emergence of the Metabolic Signaling of the Nucleoredoxin-like Genes during Evolution"

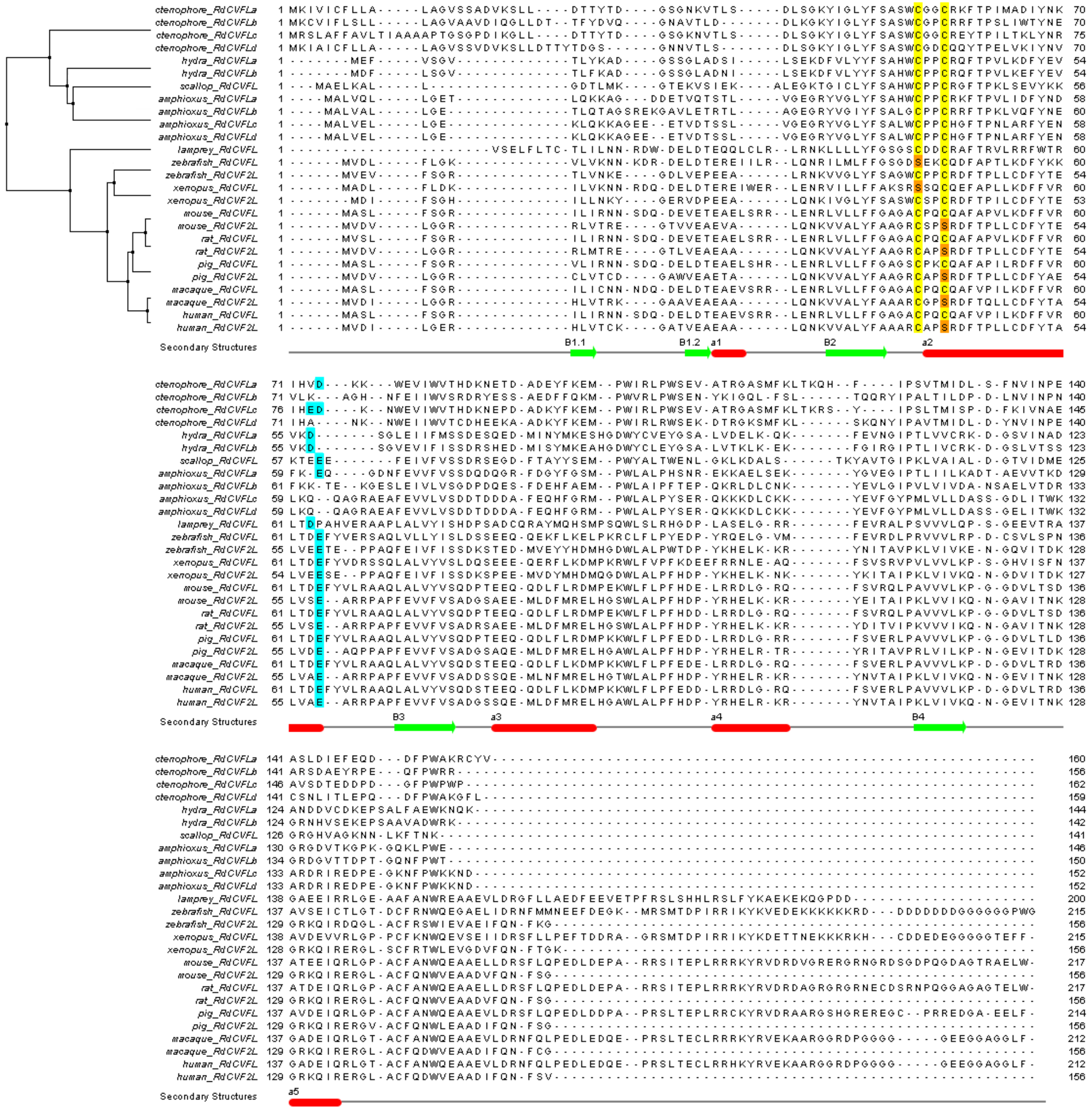


**Supplemental figure 1: Phylogenic alignment of RdCVFL proteins**

In yellow/orange, cysteine and serine in catalytic site. In blue, conserved aspartic acid or glutamic acid (E) that E plays a role in RdCVF binding to BSG1 (Aït-Ali et al, PMID: 25957687).

The beta sheet in the secondary structures are colored in green, the alpha helix in red.

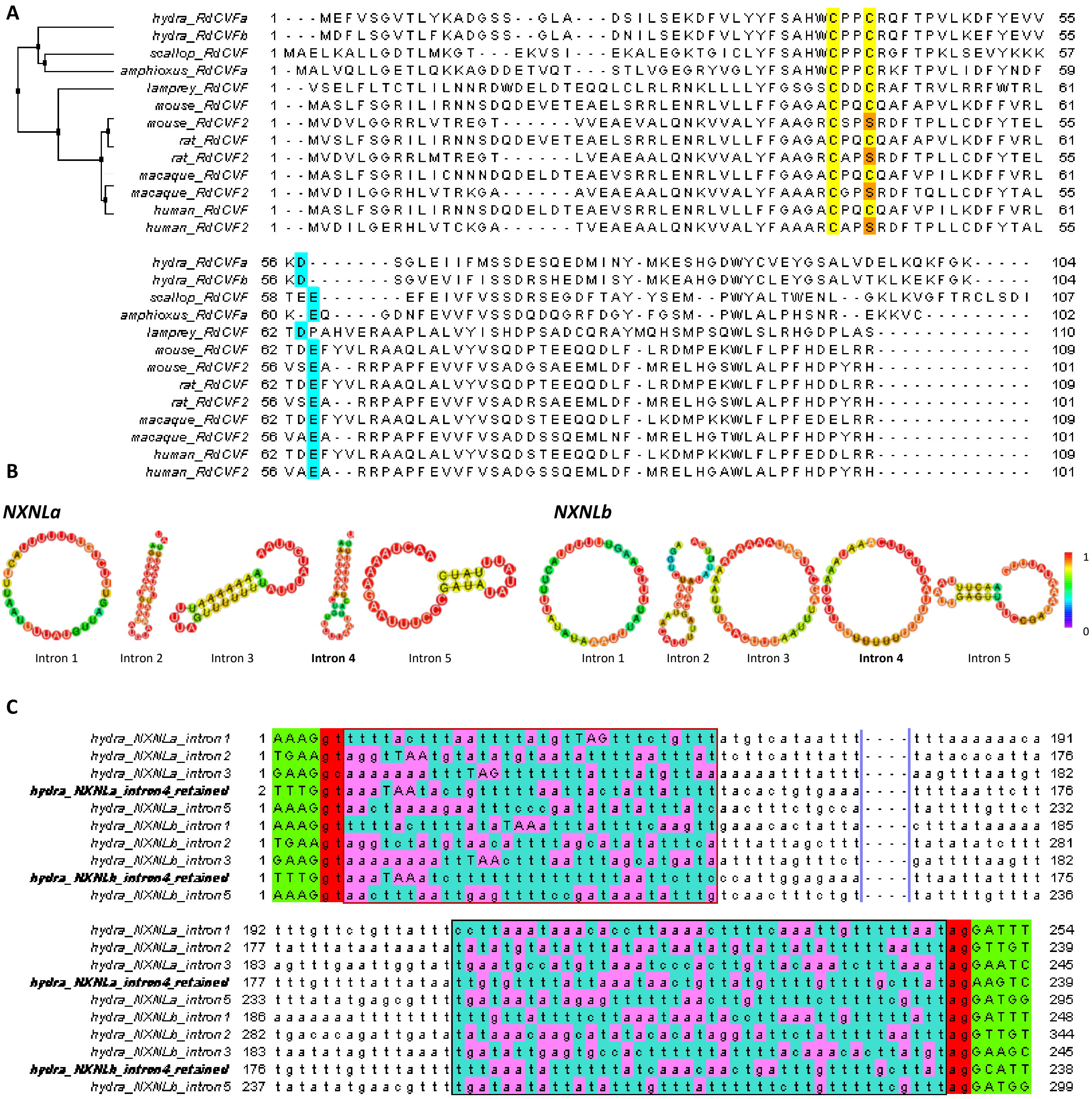


**Supplemental figure 2: Validated RdCVF proteins**

**A. Phylogenic alignment of RdCVF proteins,** In yellow/orange, cysteine and serine in catalytic site. In blue, conserved aspartic acid or glutamic acid (E) that E plays a role in RdCVF binding to BSG1 (Aït-Ali et al, PMID: 25957687).

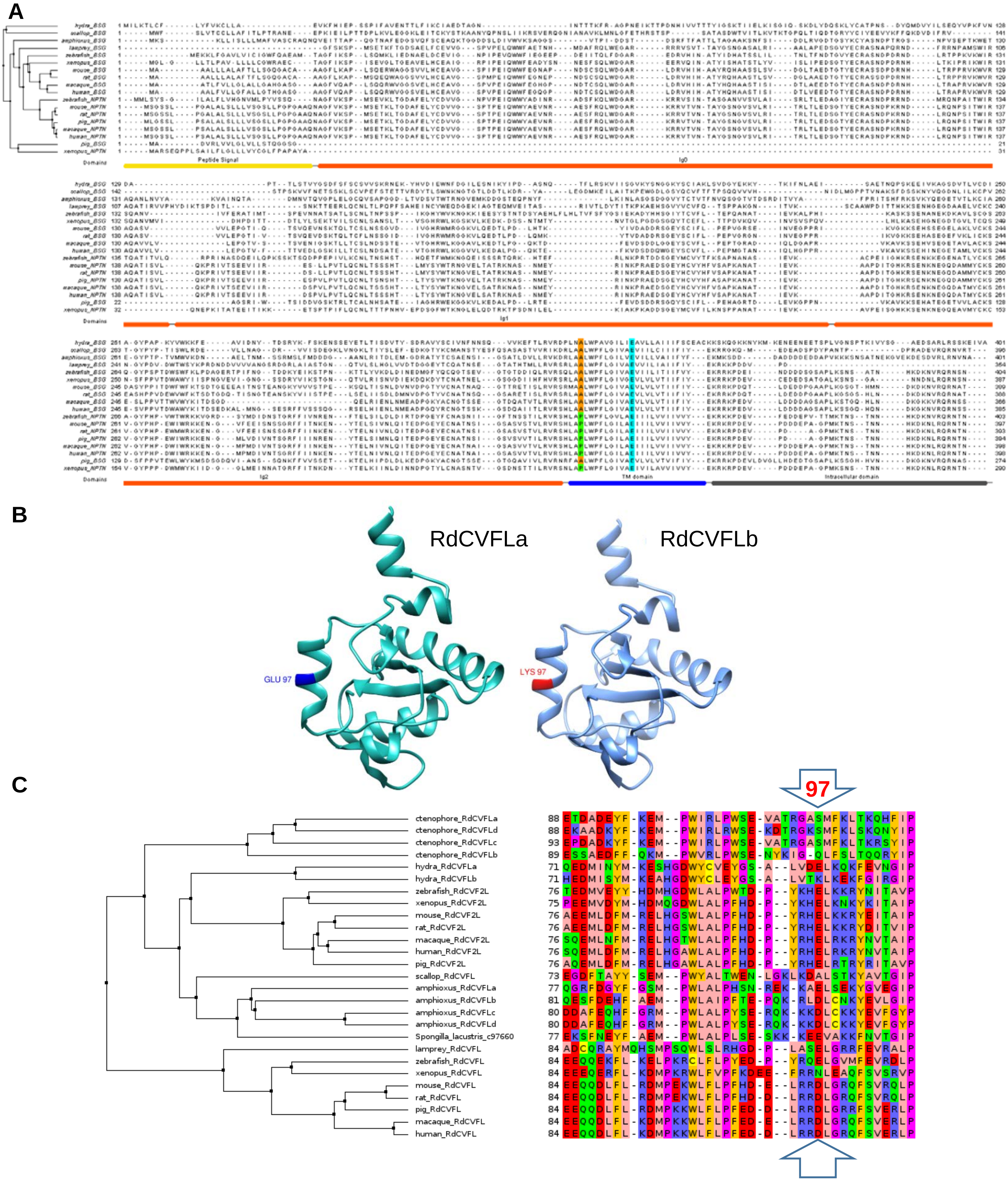


**Supplemental figure 3: Phylogenic alignment of basigin and neuroplastin proteins.** The protein domains are underlined in yellow for the signal peptide, orange for the immunoglobulin domain, blue for the transmembrane domain and grey for the intracellular domain. The colored amino acids in the highly conserved transmembrane domain are: in light blue, a conserved glutamic acid, in orange and green, conserved amino acids depending on the protein, respectively alanine for BSG and proline for NPTN.

Figure S4 full width of the page

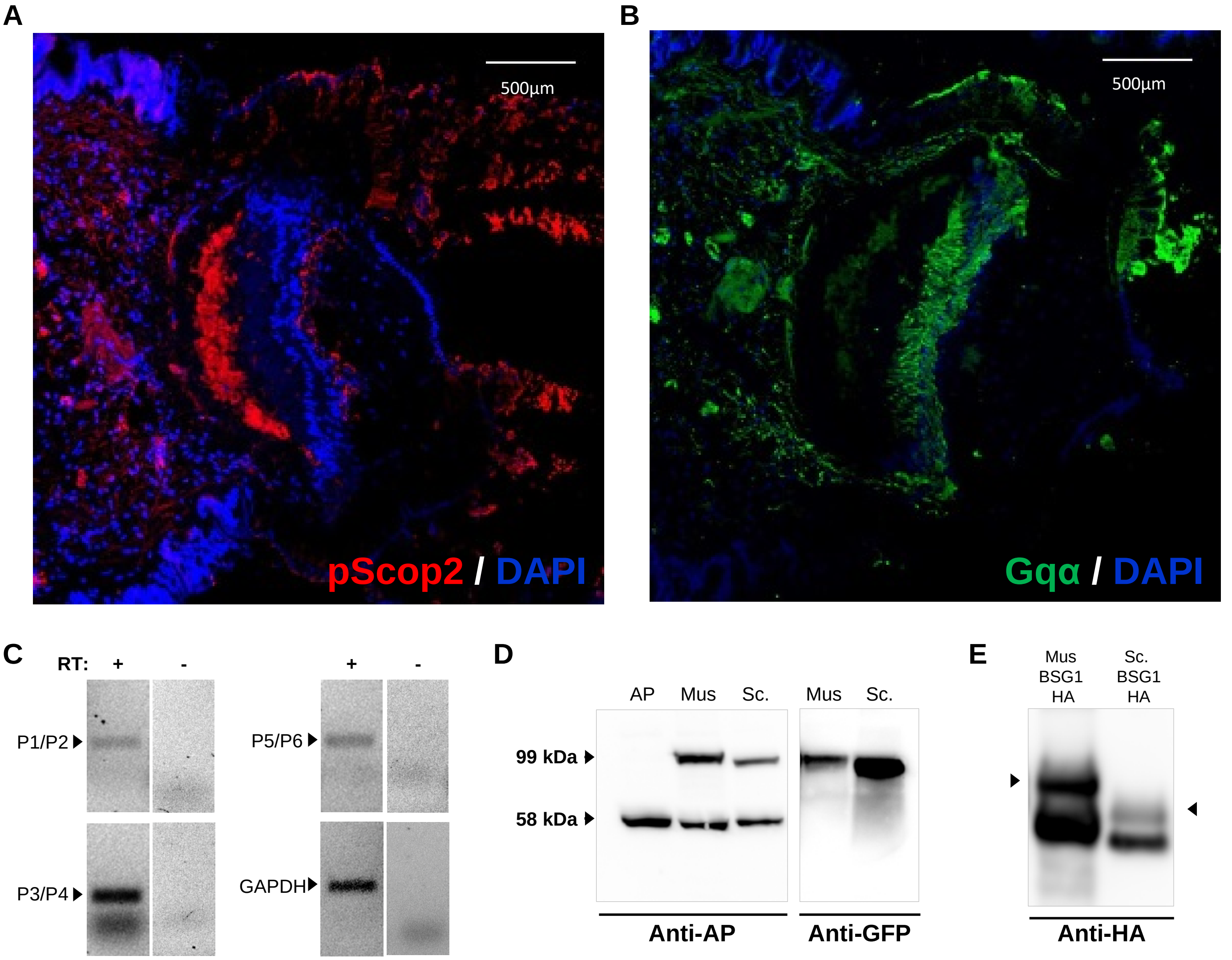

Figure S5 full width of the page

**A**

S4RAQ8: *Petromyzon marinus* basigin-1 (354 amino-acids)

Ig, Ig1 and Ig2

GFSKSPMSETKFTGDSAELFCEVVGSPVPELQWWFAETNHMDAFRQLWEG
ARRRRVSVTTAYGSNGASALRIAALAPEDSGVYECRASNAPQRNDFRRNP
AMSWIRAQATIRVVPHYDIKTSPDITLSNKTTEERLQCNLTLPQPFSAHE
INCYWEQDGEKIAGTEQMVEITASRIVTLDYTITKPKAEHSGVYVCVFQT
SPPAKGNITVKSCAAWPDITTHKKSENHGEGDAAELVCKCNGYPDVDWTW
SYKPRDNDDVVVVANGSRDGRLAIASTGNQTVLSLHGLVVDTDGGEYTCQ
ATNSEGTATHTMLLRVRSRLAALWPFLGIVAEVVILIAIIFIYEKRKKPD
DVPD

**B**

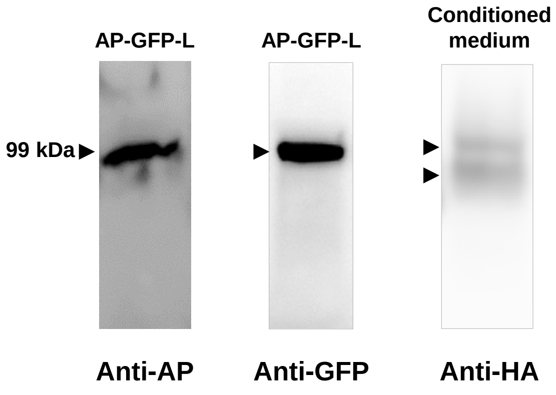

Figure S6 full width of the page

A

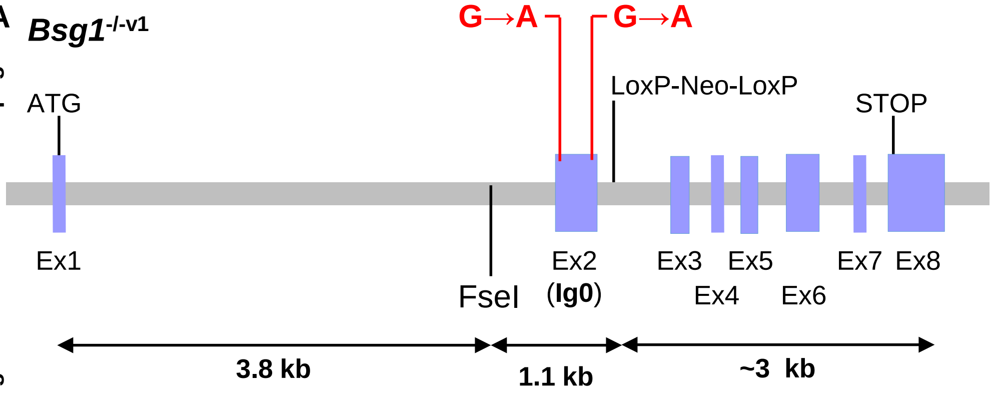


B

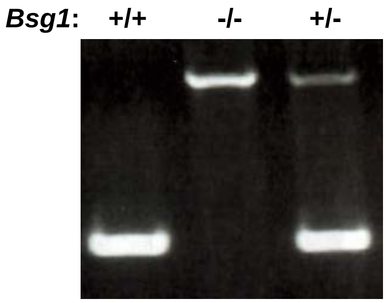


C

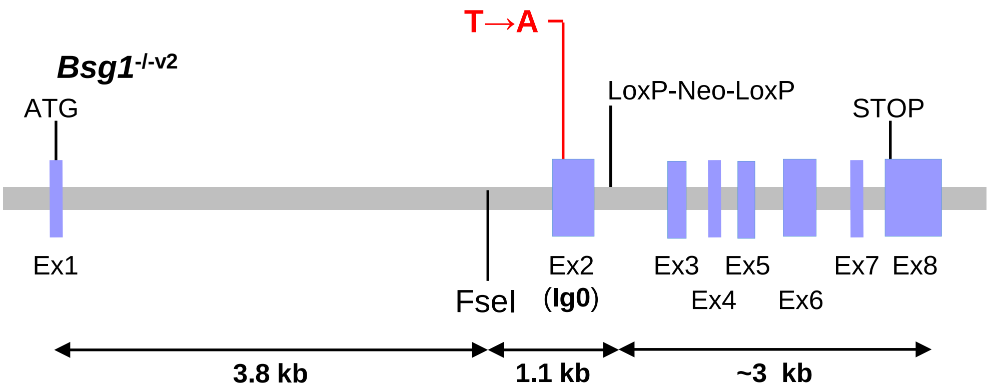


D

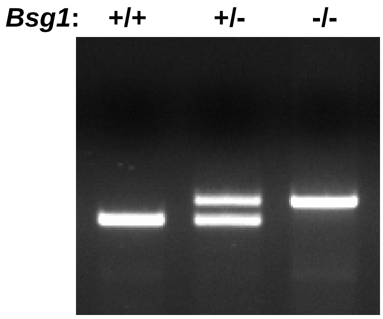


E

**Exon 2**
AY089967.1 Mus musculus basigin mRNA

CUGGUUUCCUCAAGGCACCACUGUCGCAGGAGCGGUGGGCGGGGGGCAGCG
UGA
UGGUCCUGCACUGUGAGGCUGUGGGCAGCCCCAUCCCCGAGAUCCAGUGGU
UGA
GGUUUGAAGGGAAUGCUCCAAACGACAGCUGCUCCCAGCUCUGGGAUGGUG
CCCGGCUGGACCGUGUUCACAUCCAUGCCGCCUACCGUCAGCAUGCAGCCA
GUUCGCUCUCUGUUGAUGGGCUCACCGCAGAGGACACAGGCACUUACGAGU
GCCGGGCCAGCAGUGACCCAGACCGCAACCACCUUACUCGGCCACCCAGGG
UCAAGUGGGUCCGUGCCCAGGCGAGCGUGGUGGUCCUUGAAC
UGA

G

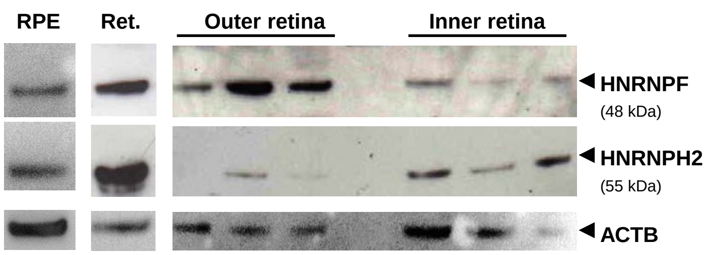


F

| Position | Splicing inhibitor | Sequence | Score *Bsg1*+/+ | Sequence | Score *Bsg1*-/-v1 | Sequence | Score *Bsg1*-/-v2 |
|---|---|---|---|---|---|---|---|
| 33-38 | HNRNP P2 (FUS) | CGGUGG | 8 | CGGUGA | 0 | CGGUGG | 8 |
| 36-40 | hnRNP H1, H2, H3 | UGGGC | 5 | UGAGC | 0 | UGGGC | 5 |
| 36-40 | hnRNP F | UGGGC | 2 | UGAGC | 0 | UGGGC | 2 |
| 41-45 | hnRNP H1, H2, H3, F | GGGGG | 6 | GGGGG | 6 | GGGGG | 6 |
| 42-46 | hnRNP H1, H2, H3, F | GGGGG | 6 | GGGGG | 6 | GGGGG | 6 |
| 43-47 | hnRNP H1, H2, H3 | GGGGC | 6 | GGGGC | 6 | GGGGC | 6 |
| 43-47 | hnRNP F | GGGGC | 4 | GGGGC | 4 | GGGGC | 4 |
| 62-66 | hnRNP A1 | CUGUG | 0 | CUGUG | 0 | CUGAG | 5 |
| 65-69 | hnRNP H1, H2 | UGAGG | 0 | UGAGG | 0 | AGAGG | 2 |
